# Decomposing scanner batch effects and their impact on pathology foundation models

**DOI:** 10.64898/2026.10.07.757122

**Authors:** Jonghyun Lee, Lina Takemaru, Yutong Zhu, William Sullivan, Matthew E. Lee, Angela Ge, Portia A. Kreiger, Amy E. Baxter, Zhi Huang, Dokyoon Kim, Derek A. Oldridge

**Author notes:** Corresponding authors. (D. Kim); (D.A. Oldridge). These authors contributed equally to this work (Jonghyun Lee, Lina Takemaru). These authors contributed equally to this work (Yutong Zhu, William Sullivan).

## Abstract

Pathology foundation models (PFMs) are expected to represent the same tissue in the same way on any scanner, but their embeddings still carry scanner signals. Scanner-related image differences and the scanner sensitivity of PFMs have each been described. How tissue-dependent changes in color and spatial frequency relate to shifts in PFM representations, and what this means for correction, is less understood. We imaged 103 slides from 37 normal tissue types on six scanners (PanNormal; 95,780 matched locations) and used the real paired images as targets. We used the public PLISM dataset for external validation. First, we decomposed scanner differences into image properties. Color and spatial frequency changed together, by amounts that depended on tissue type. Second, corrections that matched these measured properties brought images closer to the target more reliably than representations. Pix2Pix reduced the image residual by 69% but increased UNI distance to the target by 27%. The same frequency matching helped one scanner and hurt another. Probes still distinguished corrected from real target embeddings after every correction. Third, changing one frequency band at a time showed that PFMs differ in how strongly each band moves them and whether it moves them toward the target. Across eight PFMs pretrained on whole-slide image tiles, the models that moved least under these small changes were the most robust to real scanner differences (Spearman *ρ* = −0.90). Aligning images and aligning representations are therefore different goals. The gap between them depends on scanner, tissue, and model.

**Highlights:**

- Same slides on six scanners link image changes to PFM representation shifts.
- Scanner changes in color and frequency vary by tissue type.
- Corrections aligned images more than representations and never matched real targets.
- PFMs respond differently to frequency bands, so one correction can help or hurt.
- PFMs less sensitive to small frequency changes were more robust across scanners.

## 1. Introduction

Pathology images of the same tissue can look different when they come from different institutions or scanners. Color, contrast, and the sharpness of fine structure all change (Kothari et al., 2014; Swiderska-Chadaj et al., 2020; Stacke et al., 2021; Howard et al., 2021). Pathology foundation models (PFMs) are pretrained on images from many institutions and scanners. They are expected to represent the same tissue in the same way regardless of how it was scanned (Chen et al., 2024; Lu et al., 2024; Zimmermann et al., 2024; Xiang et al., 2025; Alber et al., 2025; Ding et al., 2025; Scalbert et al., 2026). In practice, PFM embeddings still carry scanner signals. They can identify the scanner or the medical center of an image (Carloni et al., 2025; Thiringer et al., 2026; de Jong et al., 2025). These signals can affect downstream predictions (Kömen et al., 2026). Recent work has therefore started to build robustness into the models themselves (Alber et al., 2026; Hemker et al., 2026).

Earlier studies have described this problem from two sides, the image and the representation. On the image side, datasets in which the same tissue was scanned on several devices showed that the resulting images differ in color, contrast, sharpness, and texture (Wilm et al., 2023; Ochi et al., 2024). Measurements with standardized test objects, whose true appearance is known, showed the same kinds of differences across nine scanner models (Pye et al., 2026). Tools for detecting batch effects summarize such differences as simple image statistics, such as intensity, gradient, and texture features, and compare them between sites (Wu et al., 2025). On the representation side, embeddings of the same slides shift when the slides are scanned on different devices. These shifts change prediction consistency (Thiringer et al., 2026). Robustness indices measure how strongly technical signals compete with biological signals (Kömen et al., 2026). Benchmarks with synthetic blur, noise, stain, and contrast changes measure how far embeddings drift (Coşar, 2026).

Two kinds of correction are used to reduce scanner signals. Image-level correction changes the image before it reaches the model, through stain normalization (Reinhard et al., 2001; Macenko et al., 2009; Vahadane et al., 2016), augmentation (Tellez et al., 2019) or learned image translation (Bentaieb and Hamarneh, 2018; Isola et al., 2017; Zhu et al., 2017). Because the corrected image can be given to any model, this approach is model-agnostic. Feature-level correction changes the embeddings of one model, for example through domain adaptation (Ganin et al., 2016; Ren et al., 2019; Lafarge et al., 2019; Kim et al., 2025), batch-effect correction (Johnson et al., 2007) or paired affine alignment as in FEATMAP (Donle et al., 2026). Both kinds of correction are usually judged by indirect measures: whether corrected images look more like the target domain, or whether embeddings from different scanners move closer together. These measures can look good even when a correction moves an image toward the wrong target or removes the tissue information that the embedding should keep.

The link between the two sides is less understood. Representation studies have measured how far embeddings move between scanners (Thiringer et al., 2026; Kömen et al., 2026) or under synthetic image changes (Coşar, 2026). They have not shown which of the measured image changes move embeddings toward or away from the real target. How do tissue-dependent changes in color and spatial frequency relate to shifts in the representations of different PFMs? What does this mean for correction? Answering this requires a real target: the same tissue location imaged on the target scanner. Imaging the same slides on several scanners provides such targets. It also keeps the tissue fixed, so the remaining differences come from acquisition.

Here we imaged 103 slides from 37 normal tissue types on six scanners (PanNormal). The paired images at 95,780 matched locations served as targets, and PLISM served as an external dataset. With these targets, we decomposed scanner batch effects into color, contrast, and spatial-frequency changes. We then combined this decomposition with controlled image changes to test how PFMs respond. We studied four widely used PFMs in depth and extended the main comparisons to eight further models. These included vision–language models, models fine-tuned with spatial-transcriptomic anchors, a stain-normalized model, and three natural-image references (Lu et al., 2024; Huang et al., 2023; Hemker et al., 2026; Yun et al., 2024; Oquab et al., 2024; Radford et al., 2021; He et al., 2016). We asked three research questions (RQs):

**RQ1.** What do scanners change in the image, and does this change depend on tissue type?

**RQ2.** When corrections match the measured image properties, do PFM representations follow?

**RQ3.** How do PFMs respond to controlled image changes, and does this response relate to their robustness to real scanner differences?

Scanner image changes depended on tissue (RQ1). Matching these changes aligned images more reliably than representations (RQ2). The strength of a PFM’s response to small frequency changes tracked its robustness to real scanner differences (RQ3).

## 2. Materials and Methods

### 2.1. Datasets and paired locations

To separate scanner effects from biological variation, we assembled PanNormal by imaging the same physical slides repeatedly on six scanners: AT2, GT450, S360, S60, VERSA, and AKOYA. PanNormal contains 103 slides from 37 normal tissue types (Supplementary Table S1). Comparing the same tissue section holds tissue composition constant across scanners. Here, scanner effect denotes the effect of the whole acquisition pipeline, including scanner model, individual device, imaging time, and vendor processing. Native resolution and acquisition magnification of each scanner are given in Supplementary Table S2.

We designated AT2 as the reference scanner and registered the other scanners to its coordinate system with VALIS rigid registration (Gatenbee et al., 2023). We selected AT2 patches with a tissue fraction of at least 0.90. We confirmed corresponding locations across scanners by local optical density (OD) gradient matching. We then resampled all images onto a common 0.5052 *μ*m/px grid. Each 256 × 256-pixel image therefore covered the same 129.3 *μ*m field of view. If any of the six scanner images failed registration or tissue-quality criteria, we excluded the whole location. To reduce imbalance due to slide size, we sampled locations evenly across 10×10 spatial bins. We kept at most 1,000 locations per slide. This gave 95,780 fully matched locations. From these, we set aside 40 locations per slide for fitting corrections and a subset of 20 per slide (2,060 locations) for evaluation. All comparisons between methods used the same 20 locations (Supplementary Methods).

As an external dataset, we used the public PLISM dataset. In PLISM, one tissue microarray was prepared as 13 serial sections. Each section was stained under a different H&E condition and imaged on seven scanners (Ochi et al., 2024). PLISM contains 46 tissue cores and shares four scanner models with PanNormal: AT2, GT450, S360, and S60. After conversion to the same field of view and grid as PanNormal, we used at most four locations per section and core. This gave 2,387 locations.

### 2.2. Pathology foundation models

We used four widely used PFMs as frozen encoders: UNI and UNI2-h (Chen et al., 2024), Virchow2 (Zimmermann et al., 2024), and H-optimus-1 (Scalbert et al., 2026). UNI was the primary model. The other three were used to test whether findings held across models. To ask why some models are more robust than others (RQ3), we added six models that differ in their training signal or data. CONCH pretrained its image encoder on whole-slide image (WSI) tiles and then aligned it with image–caption pairs from publications and educational sources (Lu et al., 2024). PLIP was trained on pathology image–text pairs shared on social media (Huang et al., 2023). SEAL-UNI2 and SEAL-CONCH fine-tune UNI2-h and CONCH with spatial transcriptomic anchors (Hemker et al., 2026). EXAONEPath was pretrained on stain-normalized WSI tiles and was used with its released Macenko normalization (Yun et al., 2024). DINOv2 ViT-L/14, pretrained on natural images, served as a non-pathology reference (Oquab et al., 2024). Two further natural-image references were CLIP ViT-B/32 (Radford et al., 2021) and ResNet50 trained on ImageNet (He et al., 2016). PLIP was fine-tuned from CLIP ViT-B/32. ResNet50 is a common feature extractor in WSI pipelines (Lu et al., 2021). CLIP and PLIP share architecture and preprocessing and differ only in their weights. Every model received the same 129.3 *μ*m image and used its released preprocessing, except that ResNet50 saw the whole field without a center crop. The released center crops of EXAONEPath and DINOv2 reduced their effective field of view to 113.2 *μ*m (Supplementary Methods).

### 2.3. Evaluation against real paired targets

Every correction mapped reference-scanner images toward one target scanner and was compared with the real target-scanner image at the same location. We judged each correction on three axes.

#### Image fidelity

We measured 10 image properties spanning color, OD, contrast, gradient energy, and three frequency bands (Section 2.4). Image residual summarized how far these properties of a corrected image were from those of its target, after scaling each property by its spread on training slides. Lower values mean closer to the target. Joint coverage was the proportion of locations at which all 10 properties fell within one scale unit of the target. We also used the structural similarity index (SSIM) and LPIPS–VGG16. SSIM compares local brightness, contrast, and structure (Wang et al., 2004). LPIPS–VGG16 is a distance between learned perceptual features (Zhang et al., 2018).

#### Representation alignment

Target distance was the cosine distance between the embedding of the reference-scanner image, raw or corrected, and the embedding of the real target-scanner image at the same location. Target gain was the reduction in this distance from raw. Positive gains mean movement toward the real target. To compare models with different distance scales, we divided target distance by the mean distance between tissue types in the same model. We call this normalized scanner distance. Detectability asked whether a classifier could still tell corrected images from real targets. We used three probes of increasing flexibility (linear, multilayer perceptron, and *k*-nearest neighbors) and report the best of the three. Chance-level accuracy is 0.5. For raw images, this measures the scanner signal. After correction, it can also reflect artifacts introduced by the correction itself. Probes were trained and tested only on slides held out from the correction fit, so that no probe saw data used to fit the correction.

#### Tissue information

We averaged embeddings over the evaluated locations of each slide. For each slide, we then found the nearest slide among real target-scanner images of other physical slides. We checked whether this nearest slide had the same tissue type. We report macro recall over the 36 tissue types represented by at least two slides.

#### Validation design

PanNormal was the primary dataset and was split into five folds by physical slide. All corrections were fitted on training folds and evaluated on held-out slides. PanNormal results summarize the five directions from the reference scanner to the other scanners. In PLISM, we evaluated the three directions shared with PanNormal (reference scanner to GT450, S360, and S60) and applied corrections fitted in PanNormal without refitting. We used PLISM to test whether the scanner phenotypes (RQ1) and the corrections (RQ2) carried over to another acquisition context. The analyses for RQ3 used PanNormal only. We report PanNormal results unless PLISM is named.

### 2.4. Scanner phenotypes and their tissue dependence (RQ1)

We decomposed the differences between images of the same tissue location into three groups: color, contrast and edges, and spatial frequency. We used CIELAB because it separates lightness (*L*^∗^) from two color directions, green to red (*a*^∗^) and blue to yellow (*b*^∗^). Mean OD summarized how dark the tissue appeared. The standard deviations of *L*^∗^ and OD, together with OD-gradient energy, measured contrast and edge strength. For spatial frequency, we divided the Fourier amplitude of OD images into low–mid (0.10–0.30), mid (0.30–0.60), and high (0.60–0.90 cycles/*μ*m) bands. Lower frequencies capture broad tissue patterns, and higher frequencies capture small edges and fine structure. For each property, the difference between paired reference-scanner and target-scanner images defined a scanner phenotype. We summarized scanners on two axes: the overall color difference from the reference scanner (CIELAB Δ*E*_76_) and the log_2_ ratio of high-band amplitude in the target image to that in the reference image. To show the full frequency response, we also computed transfer curves. A transfer curve is the log_2_ amplitude ratio to the reference scanner at each spatial frequency. We averaged the curves within tissue type (Supplementary Methods).

To test whether scanner effects depend on tissue type, we analyzed 13 image measures: the 10 above plus gradient agreement with the reference image, tissue fraction, and Laplacian variance. For each slide and target scanner, we computed the difference from the reference scanner. This gave 515 contrasts from 103 slides. For each measure, we fitted a linear mixed-effects model with a mean effect for each target scanner and random effects for tissue type, for physical slide, and for each scanner within each tissue type. The last of these, the scanner-by-tissue effect, lets the scanner effect differ between tissue types. We tested tissue dependence with likelihood-ratio tests calibrated by parametric bootstrap. We adjusted the *p* values across the 13 measures by the Benjamini–Hochberg method. After removing the scanner means, we report the share of the remaining variance explained by the scanner-by-tissue effect (Supplementary Methods).

### 2.5. Image-level correction (RQ2)

We compared four families of image-level correction, from parametric augmentation to a correction targeted at the scanner phenotypes measured for RQ1.

#### Augmentation

We built 257 augmentation candidates from the original image and random draws of color, H/E stain scaling, brightness, contrast, saturation, gamma, blur or sharpening, and noise. A smaller set of 65 candidates used a narrower range. We first selected one candidate per scanner to minimize image residual on training slides (global setting). We then selected, at each location, the candidate closest to the real target (oracle). Because the oracle uses the target at test time, it is not a usable correction. It shows the best that the augmentation set can reach.

#### Blur and sharpening trajectories

To see how augmentation moves PFM representations, we applied Gaussian blur and unsharp masking at increasing strengths to images from all six scanners (Supplementary Methods). We then followed the UNI embeddings of each scanner as strength increased. We measured whether scanners moved in the same direction, how much the distance between scanners shrank, how much fine structure remained, and whether the nearest slide still had the same tissue type. A six-way linear probe tested whether the scanner could still be identified after strong blur.

#### Stain normalization and learned translation

Stain normalization used Reinhard, Macenko, and Vahadane (Reinhard et al., 2001; Macenko et al., 2009; Vahadane et al., 2016). Image translation used paired Pix2Pix and unpaired CycleGAN (Isola et al., 2017; Zhu et al., 2017), each trained from the reference scanner to each target scanner. Pix2Pix learned from paired targets at the same locations, whereas CycleGAN sampled source and target images independently. *Color–frequency correction.* To test whether correcting both phenotype axes of RQ1 would bring images and representations closer to the target, we applied Reinhard color normalization followed by frequency matching.

Frequency matching multiplied the Fourier amplitude of OD images by scanner-specific gains estimated from paired spectra on training slides. It kept phase and color unchanged. We also applied each step alone.

### 2.6. Feature-level correction (RQ2)

We compared feature-level correction with image-level correction against the same paired targets. For each PFM and target scanner, we fitted a paired affine map with ridge regularization, as in FEATMAP (Donle et al., 2026). We also fitted ComBat with the target scanner as reference batch (Johnson et al., 2007). Maps were fitted on 40 locations per training slide. They were applied to the 20 evaluation locations of held-out slides and, without refitting, to PLISM. Least-squares affine maps are reported in Supplementary Section B.8.

### 2.7. Frequency sensitivity and scanner robustness of PFMs (RQ3)

To test how PFMs respond to spatial frequency, we changed one frequency band at a time in color-corrected reference-scanner images. For each image, we raised or lowered the amplitude of the low–mid, mid, or high band so that every change had the same OD magnitude (dose; Supplementary Methods). We then measured the cosine distance between the embeddings before and after the change. We also changed each band in the direction of the real target scanner and measured whether the embedding moved toward the target. Neither the target image nor the PFM was used to set the size of a change.

We then asked whether these responses relate to robustness to real scanner differences. For each model, frequency sensitivity was the representation shift caused by the same small change, divided by the model’s mean between-tissue distance. This division let us compare models with different distance scales. The band ratio, high-band divided by low–mid-band sensitivity, described which band a model weighted more. As the main robustness measure, we used the PathoROB robustness index (Kömen et al., 2026). For each image, the index looks at its nearest images in embedding space. Among them, it counts those showing the same tissue on another scanner and those showing another tissue on the same scanner. The index is the first count divided by the sum of both counts. A higher index means that tissue, rather than scanner, organizes the representation. We excluded neighbors from the same physical slide. With 36 tissue types and six scanners, random neighbors give an index of 0.089. Normalized scanner distance and detectability served as secondary robustness measures. Tissue retrieval showed whether a robust model still carried tissue information. Robustness and tissue-retrieval measures were computed from raw images. Detectability separated reference-scanner from target-scanner embeddings. Retrieval searched raw reference-scanner images of other slides (Supplementary Methods).

We related sensitivity to robustness by Spearman correlation across the eight models whose image encoders were pretrained on WSI tiles: UNI, UNI2-h, Virchow2, H-optimus-1, EXAONEPath, SEAL-UNI2, CONCH, and SEAL-CONCH. PLIP, CLIP, DINOv2, and ResNet50 are shown for reference. Their image encoders were trained on natural images or started from such an encoder. EXAONEPath without its normalization is also shown for reference. To place these reference models, we fitted a least-squares line across the eight models. We then asked whether the residual of each reference model fell within the range of residuals of the eight.

Finally, we asked whether the frequency weighting of a PFM depends on the scanners that produced its pretraining images. We continued the self-supervised pretraining of UNI and Virchow2 on PanNormal images from two scanners, AT2 and GT450. GT450 images have higher high-frequency amplitude than AT2 images (Section 3.1). Each training location contributed one image, from either AT2 or GT450. Only the AT2 share of these images differed between runs (11 shares from 0 to 1 for UNI and six for Virchow2). All runs started from the released weights, used the DINOv2 objective (Oquab et al., 2024) without blur augmentation, and were repeated with two data orders. After one epoch, we measured band sensitivity and the band ratio, as above, on held-out slides that were not used for pretraining (Supplementary Methods).

### 2.8. Statistical analysis

In PanNormal, we first averaged locations within each slide and scanner, weighted the five target scanners equally within each slide, and summarized over the 103 physical slides. In PLISM, we averaged within section and summarized over the 13 sections. We report means with 95% confidence intervals (CIs) from 2,000 bootstrap resamples of physical slides (PanNormal) or sections (PLISM). Paired differences used the same resamples. Percentage changes were defined so that positive values mean improvement. To summarize how image alignment related to representation alignment, we computed Spearman correlations across the 35 combinations of image correction and target scanner, separately for each PFM. For the sensitivity–robustness analysis, CIs for Spearman’s *ρ* came from recomputing *ρ* in every slide resample. The *p* values came from an exact permutation test over all orderings of the eight models. These CIs describe uncertainty from slide sampling for a fixed set of models, not from the choice of models. Further details are given in Supplementary Methods.

## 3. Results

### 3.1. Scanners change color and fine detail together, by amounts that depend on tissue (RQ1)

Earlier studies found that PFM embeddings encode the scanner and the medical center (de Jong et al., 2025; Kömen et al., 2026; Thiringer et al., 2026). PanNormal reproduced this finding. A linear probe identified the scanner from raw UNI embeddings with 99% balanced accuracy (chance 17%). Because tissue was held fixed, the paired images also showed how large this signal is. The largest mean distance between scanners, between AT2 and AKOYA, was 39% of the mean distance between different tissue types.

To see what lies behind this signal, we decomposed the paired image differences into color, contrast, and spatial-frequency changes. Raw paired examples show how the same tissue appears on all six scanners (Fig. 1). On shared axes of color distance and high-frequency amplitude, AKOYA and GT450 lay at similar color distances from AT2 (Fig. 2A). However, AKOYA images had lower high-frequency amplitude than AT2 images (log_2_ ratio −1.8). GT450 images had higher high-frequency amplitude than AT2 images (log_2_ ratio +0.7). The transfer curves show where in the spectrum these differences arose (Fig. 2B). AKOYA lost amplitude steadily from about 0.1 cycles/*μ*m upward. GT450 gained amplitude at high frequencies. VERSA, S360, and S60 separated from AT2 mainly above about 0.6 cycles/*μ*m. Scanners thus differed by distinct combinations of color and spatial frequency.

**Figure 1:**
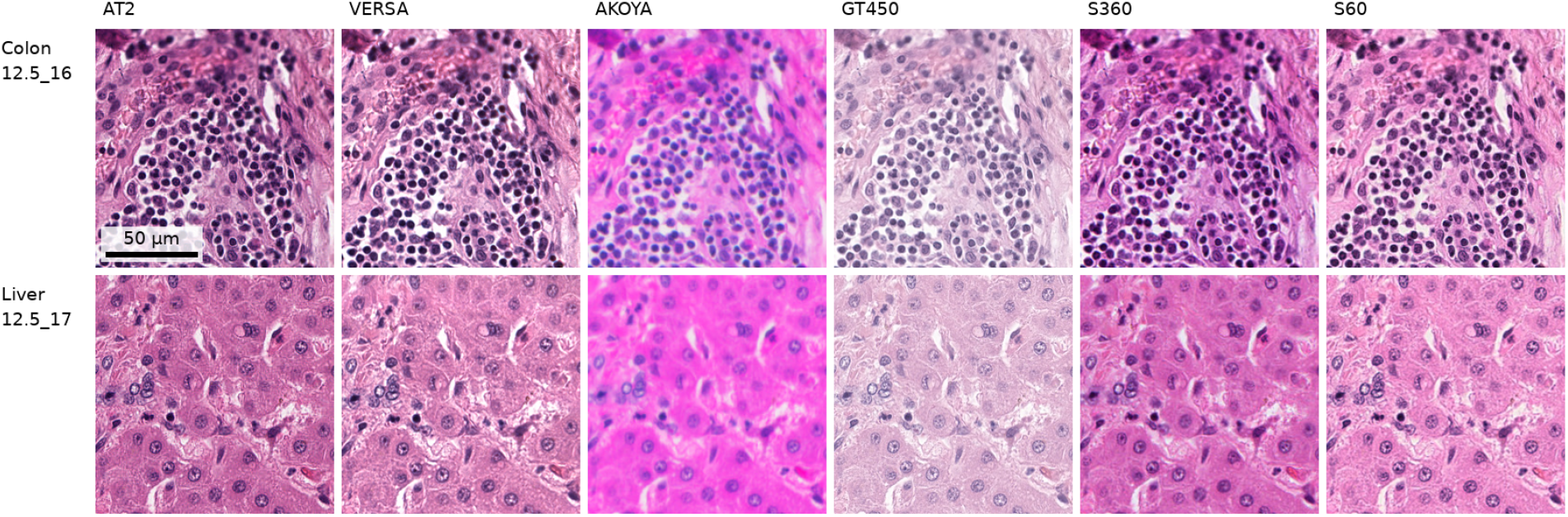
The same tissue imaged on six scanners. Each row shows one registered, uncorrected PanNormal location on a single physical slide (colon, slide 12.5_16, source index 911; liver, slide 12.5_17, source index 1659). AT2 is the reference scanner. All panels show the same 129.3 *μ*m field of view on a 0.5052 *μ*m/px grid; scale bar, 50 *μ*m. Differences in color and fine structure are visible without image correction.

**Figure 2:**
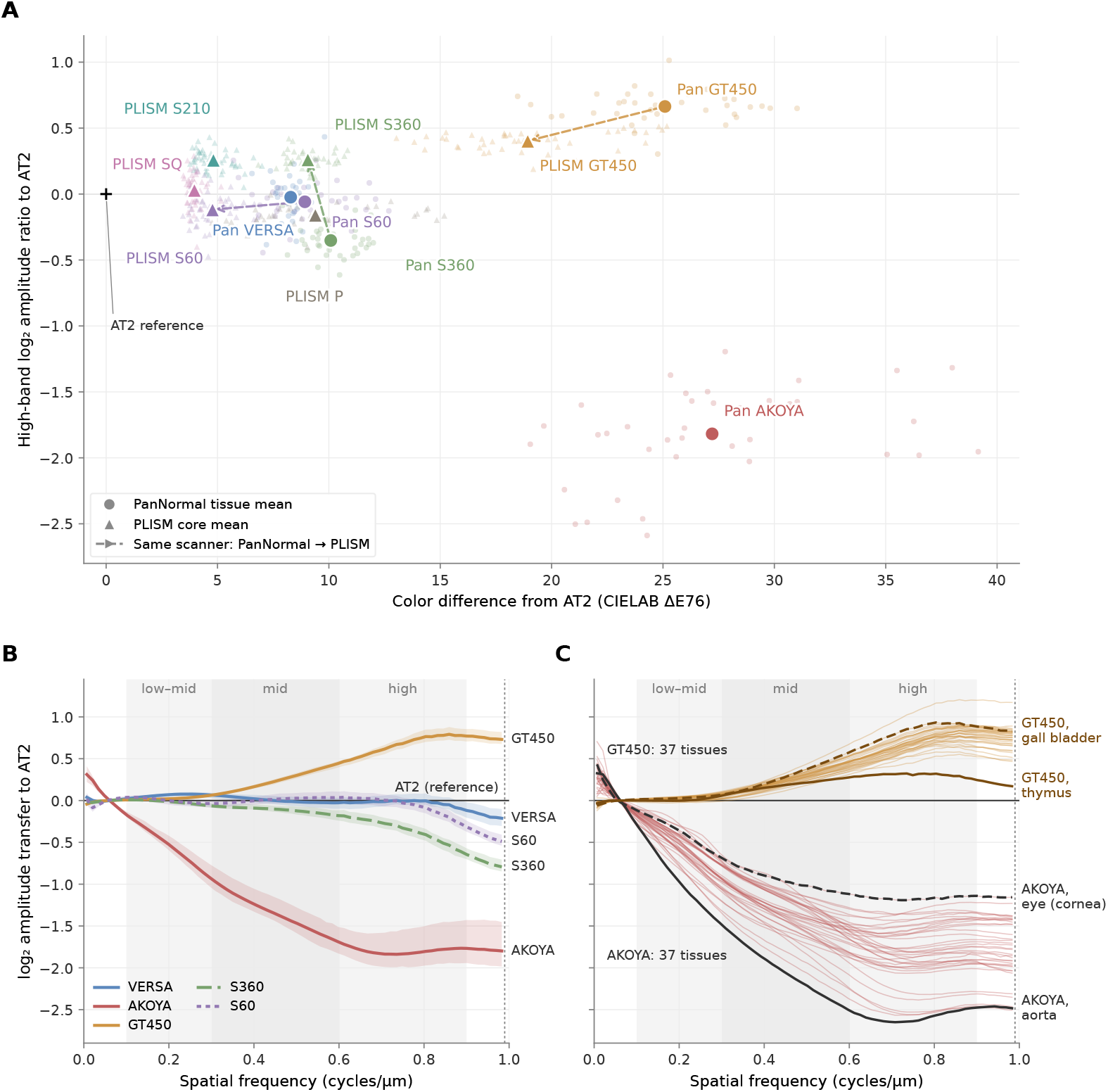
Scanners differ by distinct combinations of color and frequency. The frequency change depends on tissue. **(A)** Paired CIELAB Δ*E*_76_ from AT2 (horizontal) against the log_2_ ratio of high-band OD amplitude (vertical). Small circles show PanNormal tissue-type means. Triangles show PLISM core means. Large markers show scanner means (Pan, PanNormal). Dashed arrows connect the same scanner model in the two datasets. **(B)** log_2_ amplitude transfer to AT2 across spatial frequency, averaged over tissue types. Ribbons show the interquartile range across tissues. Shaded regions mark the three analysis bands. **(C)** Transfer curves of every tissue type for AKOYA and GT450. For each scanner, the tissue types with the lowest (solid) and highest (dashed) high-band transfer are highlighted, among those with at least two slides.

These differences depended on tissue type. In the mixed-effects model, the scanner effect differed between tissue types for all 13 image measures (Supplementary Tables S4 and S5). Tissue-specific scanner effects explained 57% of the remaining variance in low–mid-frequency transfer and 9–43% in the other measures. For example, the high-frequency log_2_ ratio of AKOYA was −2.6 in the aorta but −1.2 in the cornea. The ratio of GT450 was +0.3 in the thymus but +0.8 in the gall bladder (Fig. 2C). Tissue-by-scanner maps for all tissue types are shown in Supplementary Fig. S1. Tissue changed the size of a scanner’s effect but not the order of the most distinct scanners. In all 37 tissue types, AKOYA had the lowest high-frequency amplitude and GT450 the highest. In PLISM, GT450 again had the highest high-frequency amplitude in 44 of 46 cores. This order was also unchanged when we resampled the native scans with three different interpolation methods. In contrast, the smaller high-frequency differences of VERSA, S360, and S60 depended on the method (Supplementary Section B.2).

In answer to RQ1, scanners changed color and spatial frequency together by tissue-dependent amounts, so a correction has to change several image properties at once.

### 3.2. Matching image properties aligned images more than representations (RQ2)

RQ1 showed which image properties scanners change. We next asked whether matching these properties brings PFM representations to the real target. The image-level corrections were augmentation, stain normalization, learned translation, and a correction targeted at color and frequency. We compared them with feature-level correction.

#### 3.2.1. Augmentation did not reproduce real scanner images

We began with the simplest image-level approach: parametric augmentation of color, stain, brightness, contrast, saturation, gamma, blur or sharpening, and noise. In PanNormal, when one global candidate was chosen per scanner, the original image was selected for every scanner in all five folds. No single global augmentation brought images closer to the target than the unchanged image. We therefore selected the best candidate at each location using the real target, as an upper bound. This oracle improved SSIM and LPIPS, but it moved UNI slightly away from the target (Supplementary Table S6).

Even the oracle did not reproduce the target images. With 257 candidates, the oracle nearly halved the image residual. Yet all 10 image properties matched the target at only 49% of locations. A classifier also told real targets from oracle images with 94% accuracy.

#### 3.2.2. Blur brought scanners closer by erasing tissue information

To see how augmentation moves UNI representations, we applied blur and sharpening one factor at a time to images from all six scanners (Supplementary Fig. S3). As strength increased, the scanners moved along shared directions. The median cosine similarity between their displacement directions was 0.77 for blur and 0.70 for sharpening.

Strong blur brought the scanners together, but at the cost of tissue information. At the strongest blur, the UNI distance between scanners fell to 55% of its original value. Almost no fine structure remained. The share of nearest slides with the same tissue type fell from 64% to 45%. The scanner could still be identified with 97% accuracy. Distance between scanners can therefore shrink while the scanner remains detectable. We judged each of the following corrections by its fidelity to the real target image, its representation alignment with the real target, and its preservation of tissue information.

#### 3.2.3. Image corrections improved images more than representations

We next compared stain normalization and learned translation against real paired targets (Table 1). In PanNormal, Reinhard improved SSIM, LPIPS, and UNI distance, whereas Macenko and Vahadane increased UNI distance. Learned translation showed the clearest gap between image alignment and representation alignment. Pix2Pix reduced image residual by 69% and matched all 10 image properties at 75% of locations, yet it increased UNI distance by 27%. CycleGAN improved all three image-fidelity measures but reduced UNI distance by only 6%. The same gap appears at single locations. The correction that came closest to the target image did not come closest to the target representation (Fig. 3).

**Figure 3:**
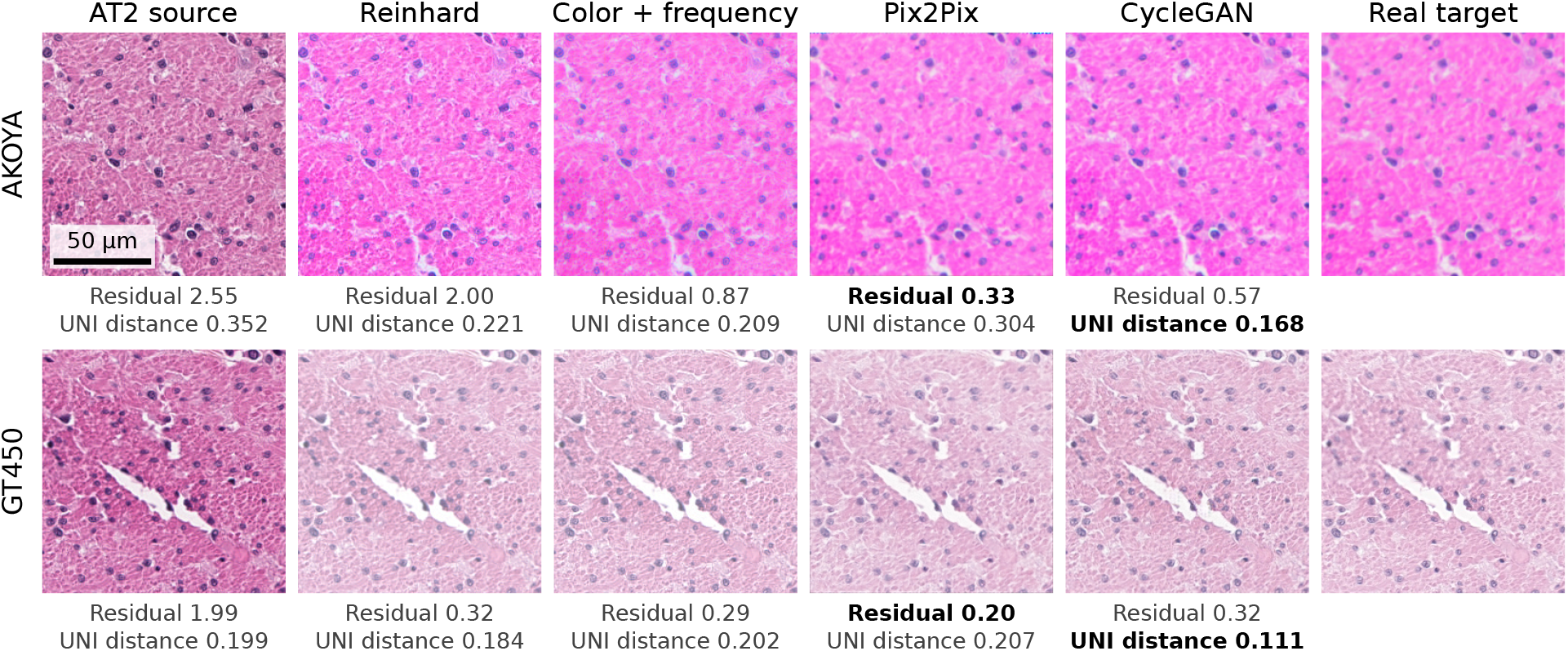
The correction with the lowest image residual does not always give the best representation alignment. Each row shows one illustrative PanNormal evaluation location (colon, slide 12.5_16) corrected from the reference scanner (AT2) toward AKOYA or GT450. Correction parameters and translation models came from the fold in which the slide was held out. Numbers give the image residual over the 10 image properties and the UNI cosine distance to the real target-scanner image. Bold marks the lowest value of each among the corrections. In both rows, Pix2Pix came closest to the target image. CycleGAN came closest to the target representation. The means of these two directions show the same pattern. Location selection is described in Supplementary Methods. All panels show the same 129.3 *μ*m field of view.

**Table 1.** Image-level correction judged on three axes against real paired target images in PanNormal and PLISM (UNI). Image fidelity: SSIM, LPIPS–VGG16, and image residual over 10 image properties (PanNormal only). Representation: UNI cosine distance to the target, its change from raw (positive = closer), and best-probe accuracy separating corrected from real target embeddings (Det.; 0.5 = chance). Tissue: same-tissue retrieval against real target-scanner images of other slides (macro recall). In PLISM, the same cores recur across sections. Retrieval is therefore near its ceiling. PLISM applies the corrections fitted in PanNormal without refitting.

| Model class | Method | Image fidelity |  |  | Representation |  |  | Tissue |
| --- | --- | --- | --- | --- | --- | --- | --- | --- |
| | | SSIM ↑ | LPIPS ↓ | Residual ↓ | Distance ↓ | $\Delta$ Raw (%) ↑ | Det. ↓ | Retrieval (%) ↑ |
| PanNormal |  |  |  |  |  |  |  |  |
| Baseline | Raw | 0.572 | 0.232 | 1.31 | 0.2136 | Ref. | 0.98 | 64.9 |
| Stain normalization | Reinhard | 0.620 | 0.202 | 0.70 | 0.1798 | +15.8 | 0.96 | 64.8 |
|  | Macenko | 0.552 | 0.244 | 1.57 | 0.3402 | -59.3 | 1.00 | 56.7 |
|  | Vahadane | 0.555 | 0.239 | 1.52 | 0.3152 | -47.5 | 1.00 | 59.6 |
| Learned translation | Pix2Pix | 0.682 | 0.218 | 0.40 | 0.2705 | -26.6 | 1.00 | 60.9 |
|  | CycleGAN | 0.658 | 0.206 | 0.55 | 0.2016 | +5.6 | 0.92 | 61.3 |
| Phenotype-targeted | Color + frequency | 0.626 | 0.192 | 0.50 | 0.1794 | +16.0 | 0.96 | 64.0 |
| PLISM |  |  |  |  |  |  |  |  |
| Baseline | Raw | 0.781 | 0.161 | - | 0.1713 | Ref. | 1.00 | 94.6 |
| Stain normalization | Reinhard | 0.817 | 0.153 | - | 0.1754 | -2.4 | 1.00 | 94.3 |
|  | Macenko | 0.775 | 0.183 | - | 0.3073 | -79.4 | 1.00 | 95.7 |
|  | Vahadane | 0.695 | 0.251 | - | 0.4150 | -142.3 | 1.00 | 93.2 |
| Learned translation | Pix2Pix | 0.699 | 0.245 | - | 0.2536 | -48.1 | 1.00 | 93.7 |
|  | CycleGAN | 0.760 | 0.234 | - | 0.2881 | -68.2 | 1.00 | 93.6 |
| Phenotype-targeted | Color + frequency | 0.809 | 0.156 | - | 0.1824 | -6.5 | 1.00 | 94.3 |

After every image correction, probes still distinguished corrected from real target embeddings. The best-probe accuracy was 0.92 or higher, compared with 0.98 for raw images. Reinhard and color–frequency correction kept tissue retrieval, but Macenko, Vahadane, Pix2Pix, and CycleGAN lowered it by 4 to 8 percentage points. A tissue classifier trained on real target images showed the same loss. For example, its accuracy for Virchow2 fell from 52% to 44% after Pix2Pix. In contrast, it classified raw reference-scanner images as well as the real targets (Supplementary Section B.7). In PLISM, every image correction moved UNI away from the target. Best-probe accuracy stayed at 0.99 or higher.

#### 3.2.4. Color–frequency correction helped AKOYA and hurt GT450

RQ1 identified color and high-frequency amplitude as the axes that separate scanners. We therefore tested a correction targeted at exactly these axes: Reinhard color normalization followed by scanner-specific frequency matching. In PanNormal, adding frequency matching reduced image residual from 0.70 to 0.50 and further improved SSIM and LPIPS. However, the mean additional UNI gain across five scanners was close to zero (Fig. 4).

**Figure 4:**
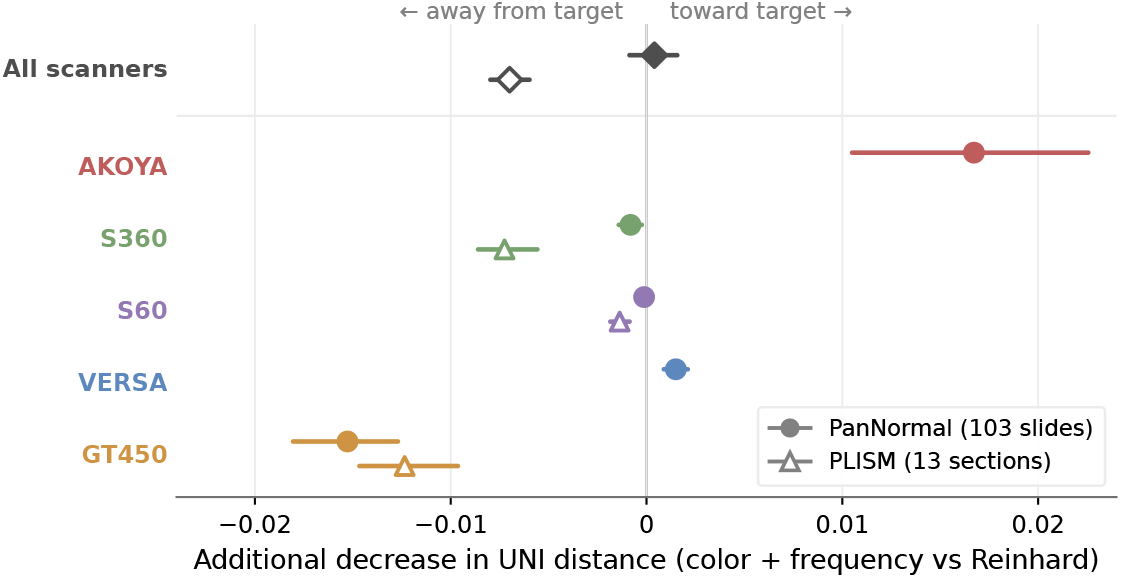
Adding frequency matching to Reinhard moves UNI toward the target for AKOYA but away from it for GT450. AKOYA has the lowest high-frequency amplitude. GT450 has the highest. The horizontal axis is the additional reduction in UNI distance to the real target-scanner image relative to Reinhard alone. Positive values indicate improvement. Rows are ordered by the PanNormal high-band amplitude ratio to AT2 (Fig. 2A). Points and lines show means and bootstrap 95% CIs across physical slides (PanNormal) or sections (PLISM). “All scanners” denotes the equally weighted scanner mean in each dataset. PLISM does not include AKOYA or VERSA.

This near-zero mean hid opposite responses that follow the scanners’ positions in Fig. 2. Frequency matching moved UNI toward the target for AKOYA, the scanner with the lowest high-frequency amplitude. It moved UNI away by a similar amount for GT450, whose high-frequency amplitude exceeded that of AT2. In PLISM, the combined correction moved UNI slightly away from the target on average.

The gains also depended on tissue. Across tissue types, the AKOYA gain from frequency matching was larger where the high-frequency loss was stronger (Spearman *ρ* = −0.57). More generally, we applied the mixed-effects model of RQ1 to correction gains. In all four PFMs, the gains of Reinhard, color–frequency, ridge, and ComBat correction differed between tissue types, and so did the added gain of frequency matching (Supplementary Section B.6). A correction tuned to the average scanner difference therefore fits some tissues better than others.

#### 3.2.5. Feature correction aligned representations within PanNormal but transferred inconsistently

Correcting features directly does not depend on how each PFM responds to image changes. Within PanNormal, ridge affine correction reduced UNI target distance by 34%. This was about twice the 16% of color–frequency correction (Table 2). Ridge affine correction also kept tissue retrieval at the level of real target images (Supplementary Section B.8). Ridge and ComBat reduced target distance in all four PFMs.

**Table 2.** Image- and feature-level correction in four PFMs, in PanNormal and PLISM. Δ: change in cosine distance to the real paired target from raw (%; positive = closer). Det.: best-probe accuracy separating corrected from real target embeddings (Section 2.3; 0.5 = chance). PLISM applies the corrections fitted in PanNormal without refitting.

|  |  | UNI |  | UNI2-h |  | Virchow2 |  | H-optimus-1 |  |
| --- | --- | --- | --- | --- | --- | --- | --- | --- | --- |
| Correction | Method | $\Delta$ (%) ↑ | Det. ↓ | $\Delta$ (%) ↑ | Det. ↓ | $\Delta$ (%) ↑ | Det. ↓ | $\Delta$ (%) ↑ | Det. ↓ |
| PanNormal |  |  |  |  |  |  |  |  |  |
| Baseline | Raw | Ref. | 0.98 | Ref. | 0.99 | Ref. | 0.96 | Ref. | 0.98 |
| Image | Reinhard | +15.8 | 0.96 | +11.4 | 0.99 | +16.4 | 0.96 | +6.5 | 0.97 |
|  | Color + frequency | +16.0 | 0.96 | +16.7 | 0.99 | +21.3 | 0.95 | +5.9 | 0.97 |
|  | Pix2Pix | −26.6 | 1.00 | +6.6 | 0.99 | −155.4 | 1.00 | −33.5 | 0.95 |
|  | CycleGAN | +5.6 | 0.92 | +32.3 | 0.93 | −30.7 | 0.93 | −12.9 | 0.94 |
| Feature | Ridge affine | +34.3 | 0.87 | +36.5 | 0.76 | +19.8 | 0.70 | +21.9 | 0.75 |
|  | ComBat | +27.7 | 0.73 | +21.0 | 0.80 | +26.8 | 0.62 | +25.1 | 0.70 |
| PLISM |  |  |  |  |  |  |  |  |  |
| Baseline | Raw | Ref. | 1.00 | Ref. | 0.99 | Ref. | 1.00 | Ref. | 1.00 |
| Image | Reinhard | −2.4 | 1.00 | +12.0 | 1.00 | −1.4 | 1.00 | +0.4 | 1.00 |
|  | Color + frequency | −6.5 | 1.00 | +9.1 | 1.00 | −3.0 | 1.00 | 0.0 | 1.00 |
|  | Pix2Pix | −48.1 | 1.00 | −33.7 | 1.00 | −191.4 | 1.00 | −64.4 | 1.00 |
|  | CycleGAN | −68.2 | 1.00 | −36.4 | 1.00 | −161.8 | 0.99 | −105.7 | 1.00 |
| Feature | Ridge affine | −36.2 | 1.00 | −38.7 | 1.00 | −67.1 | 1.00 | −57.3 | 1.00 |
|  | ComBat | −8.5 | 1.00 | +4.7 | 1.00 | +5.5 | 1.00 | −0.9 | 1.00 |

Probes also distinguished corrected from real target embeddings after feature correction. The probe that did so depended on the correction (Supplementary Table S8). After ComBat, a linear probe was close to chance (0.55–0.60), but a *k*-nearest-neighbor probe still separated the embeddings (0.62–0.80). After ridge, the pattern reversed. The linear probe separated them, while the nearest-neighbor probe was close to chance.

The internal gains transferred inconsistently to PLISM. Ridge maps fitted in PanNormal and applied without refitting increased target distance in all four PFMs. The increase was 36% in UNI. ComBat moved UNI2-h and Virchow2 slightly toward the target but UNI away from it. Best-probe accuracy stayed at 0.99 or higher for every method (Table 2).

Taken together, image alignment explained only part of representation alignment. Across image corrections and target scanners in PanNormal, the reduction in image residual was moderately related to the representation gain in UNI, UNI2-h, and H-optimus-1 (Spearman *ρ* 0.52–0.68). This relation did not hold in Virchow2 (*ρ* = 0.17). Matching color and frequency also removed only part of the representation shift. Across the four PFMs, Reinhard removed 6–16% of the raw distance to the target. Frequency matching added at most about 5 percentage points (Table 2).

In answer to RQ2, matching the measured image properties brought images closer to the target more reliably than representations. Whether representations followed depended on the scanner, the tissue, and the dataset.

### 3.3. PFM responses to small frequency changes differ between models and track robustness (RQ3)

In RQ2, the same correction moved representations toward the target for one scanner or PFM and away from it for another. To see why, we changed one image property at a time and measured how each PFM responded.

#### 3.3.1. PFMs weight frequency bands differently

We changed one frequency band at a time, with the same OD change, in color-corrected reference-scanner images (Fig. 5). The four PFMs differed in which band they weighted. UNI and UNI2-h responded more to the higher bands than to the low–mid band, whereas Virchow2 and H-optimus-1 responded most to the low–mid band.

**Figure 5:**
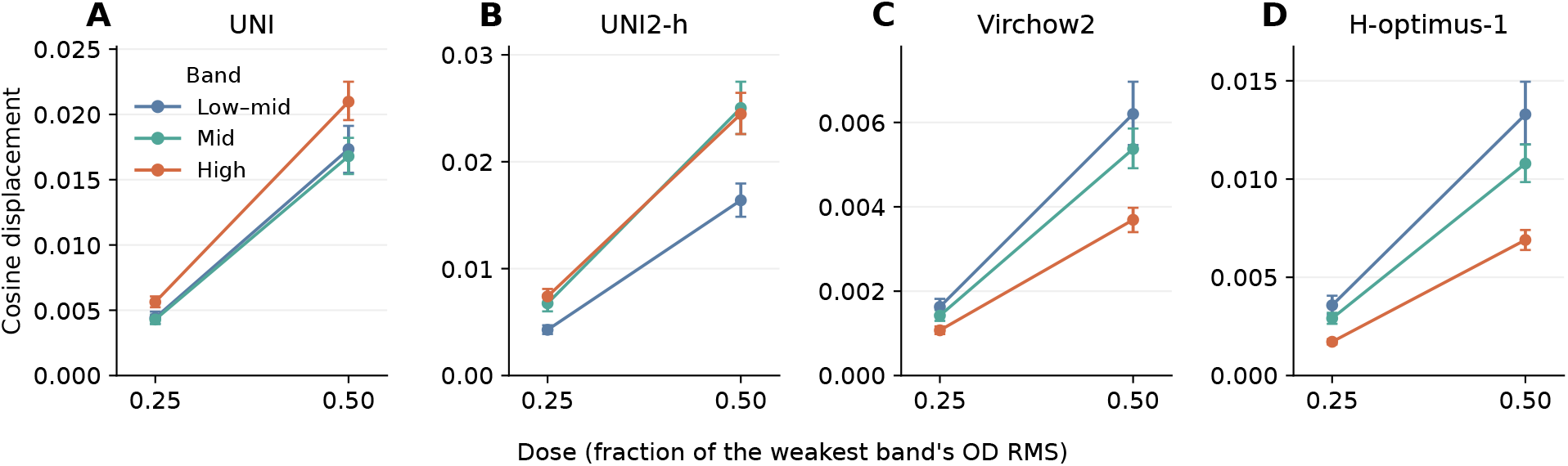
PFMs weight frequency bands differently. The low–mid, mid, or high band of color-corrected reference-scanner images was raised or lowered by the same OD magnitude (doses 0.25 and 0.50). Panels show the resulting cosine displacement of **(A)** UNI, **(B)** UNI2-h, **(C)** Virchow2, and **(D)** H-optimus-1 embeddings. Vertical scales differ between PFMs and start at zero. Points and error bars show means and bootstrap 95% CIs across 103 PanNormal slides. Each slide contributed 20 matched locations and all five target scanners.

A larger response to a band did not mean movement toward the target. When only the high band was changed in the direction of the real target scanner, UNI moved slightly away from the target on average. UNI2-h moved toward it (Supplementary Fig. S4).

#### 3.3.2. The same corrected image moves PFMs in opposite directions

Consistent with these model-specific responses, the same corrected image shifted PFMs differently (Table 2). Reinhard reduced target distance in all four PFMs. In contrast, the same CycleGAN images reduced target distance by 32% in UNI2-h but increased it by 31% in Virchow2. Adding frequency matching to Reinhard helped UNI2-h and Virchow2 in PanNormal, but not in PLISM. Across PanNormal, in 17 of 35 combinations of image correction and target scanner, the same corrected images moved at least one PFM toward the target and another away from it. An image correction can be passed to any encoder, but its effect on representation alignment was specific to the encoder.

#### 3.3.3. Anchors, heterogeneous data, and stain normalization left the scanner detectable

We next asked whether training choices explain scanner robustness. We compared the four PFMs with six further models trained with text or spatial-transcriptomic anchors, on heterogeneous image sources, or with stain normalization (Supplementary Table S10). In each of these ten models, a probe detected the target scanner with at least 95% accuracy. However, robustness varied widely, from a robustness index of 0.06 for PLIP to 0.53 for Virchow2 (chance 0.09). Two further natural-image references, CLIP and ResNet50, are added in Section 3.3.4.

These training choices did not remove the scanner signal. Their effects also differed between robustness measures. Fine-tuning UNI2-h with spatial-transcriptomic anchors (SEAL) left its detectability, scanner distance, and high-band sensitivity nearly unchanged. It did raise the robustness index from 0.30 to 0.39, because more of its nearest neighbors were the same tissue from another scanner. In CONCH, SEAL left the robustness index unchanged. CONCH, PLIP, and DINOv2 were last trained on images other than WSI tiles: image–caption pairs for CONCH and PLIP, and natural images for DINOv2. Their normalized scanner distance was not lower than that of the seven other models. They also had a lower robustness index and carried less tissue information. EXAONEPath, pretrained and applied with stain normalization, had the largest normalized scanner distance of all models and a low robustness index (0.18). Apart from the before–after comparison of SEAL, these comparisons are observational.

#### 3.3.4. Less sensitive PFMs were more robust to real scanner differences

Robustness followed how much each model moved under a small frequency change in PanNormal images (Fig. 6). Across the eight models pretrained on WSI tiles, high-frequency sensitivity was strongly related to the robustness index (*ρ* = −0.90) and to normalized scanner distance (*ρ* = +0.90). Virchow2 and H-optimus-1 moved least and were the most robust, whereas EXAONEPath moved most and was the least robust. The relation held for the mid and low–mid bands, at a larger change, after accounting for each model’s between-tissue distance, and when only one model per shared backbone was kept (Supplementary Section B.10).

**Figure 6:**
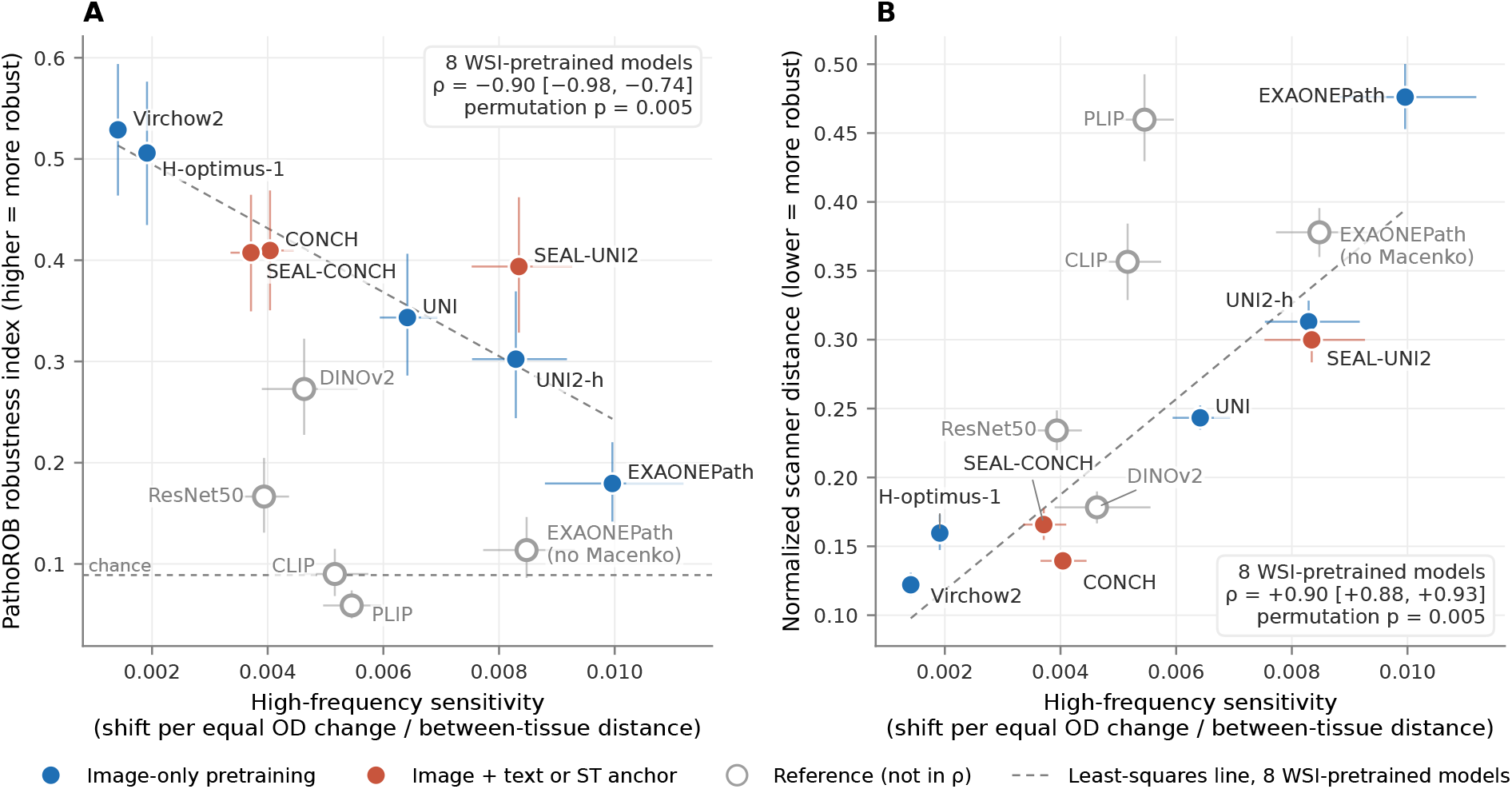
PFMs that move less under a small frequency change are more robust to real scanner differences. **(A)** High-frequency sensitivity (representation shift under an equal OD change, divided by the model’s between-tissue distance) against the PathoROB robustness index. The dashed line marks chance. **(B)** The same sensitivity against normalized scanner distance. Filled markers show the eight models pretrained on WSI tiles (blue, image only; red, with text or spatial-transcriptomic anchors). These models enter the correlations. Open markers show reference models, including CLIP ViT-B/32 and ResNet50 trained on natural images. The dashed line is the least-squares fit across the eight models pretrained on WSI tiles. Error bars and the CI of *ρ* show bootstrap 95% CIs across 103 PanNormal slides. They describe slide sampling for this fixed set of models, not the choice of models.

Robustness tracked the size of the response, not which band a model preferred. The ratio of high- to low–mid-band sensitivity was unrelated to robustness (*ρ* = −0.19; Supplementary Section B.10). Detectability was near its ceiling in every model and did not separate them.

Models whose image encoders were trained on natural images fell below this trend (Fig. 6A). PLIP, its parent model CLIP, DINOv2, and ResNet50 had moderate sensitivity, but each had a lower robustness index than the line across the eight models pretrained on WSI tiles predicted. PLIP had the lowest index. Most of its nearest neighbors were other tissues from the same scanner (Supplementary Fig. S5). CLIP showed the same pattern (robustness index 0.09, vs 0.06 for PLIP; 58% vs 63% of neighbors were other tissues from the same scanner). Most of this pattern was therefore present before pathology fine-tuning. ResNet50 had a low index for another reason. It carried little tissue information (tissue retrieval 23%) and did not have many same-scanner neighbors (Supplementary Section B.10). Across all 12 models, sensitivity still tracked normalized scanner distance (*ρ* = +0.83), but its relation with the robustness index was weaker (*ρ* = −0.51). SEAL-UNI2 also matched UNI2-h in sensitivity but not in robustness index (Section 3.3.3). The robustness index therefore reflects whether nearest neighbors share tissue type, as well as how far scanner differences move a representation.

#### 3.3.5. The scanners in the pretraining data changed frequency weighting

Published PFMs differ in architecture, data, and objective at once. Their frequency weighting cannot be traced to one of these choices. We therefore changed only the scanners in the pretraining data. We continued pretraining UNI on images of the same tissue locations. As the share of AT2 images rose, the band ratio of UNI fell steadily, from 1.38 with GT450 images only to 1.28 with AT2 images only (Fig. 7A; Spearman *ρ* = −0.99 across 22 runs from two data orders). The ratio fell mainly because low–mid-band sensitivity rose. Compared with pretraining on GT450 images only, pretraining on AT2 images only made UNI 5.0% more sensitive to low–mid-band changes and 2.1% less sensitive to high-band changes (Fig. 7B). Pretraining on the scanner with lower high-frequency amplitude thus shifted the weighting of UNI toward lower frequencies. A larger AT2 share also increased the normalized scanner distance from AT2 to the five other scanners (Supplementary Section B.11). Continued pretraining itself raised normalized scanner distance and lowered tissue classification in every run, whatever the share. We therefore compared runs with each other rather than with the released model.

**Figure 7:**
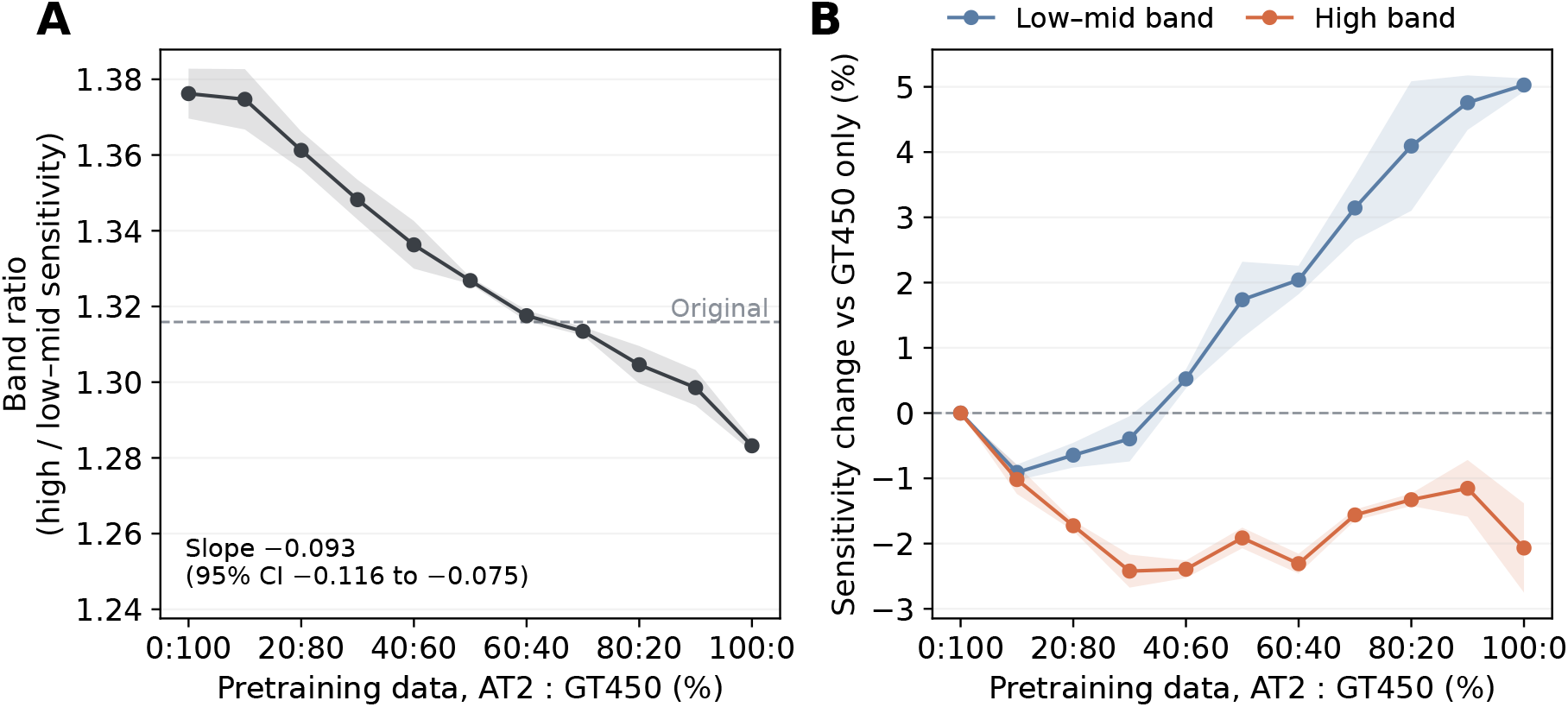
The scanners in the pretraining data shift the frequency weighting of UNI. UNI was pretrained for one more epoch on AT2 and GT450 images of the same PanNormal locations, mixed in different proportions. **(A)** Ratio of high- to low–mid-band sensitivity. The dashed line marks the original UNI. **(B)** Change in sensitivity relative to pretraining on GT450 images only. Lines show the mean of two data orders. Shading shows their range. All values are from 21 held-out slides.

The form of this change depended on the model. In Virchow2, AT2 images raised sensitivity to both bands by similar amounts (9.1% for the low–mid and 7.9% for the high band with AT2 images only). Its band ratio therefore did not change (Supplementary Fig. S6). Its normalized scanner distance increased with the AT2 share as in UNI.

In answer to RQ3, PFMs differed in how strongly and in which direction band changes moved them. Among models pretrained on WSI tiles, the overall size of this response tracked robustness to real scanner differences. Changing the scanners in the pretraining data changed the response differently in UNI and Virchow2.

## 4. Discussion

We decomposed scanner batch effects into measurable image changes and linked them to how PFMs represent the same tissue. Earlier studies described scanner differences with summary statistics of images, test objects, or repeated scans (Stacke et al., 2021; Wilm et al., 2023; Ochi et al., 2024; Pye et al., 2026; Wu et al., 2025). Paired images of the same tissue allowed us to describe each scanner as an imaging system. The frequency transfer curves measure spectral transfer on real tissue, relative to the reference scanner. They show at which spatial frequencies each scanner gains or loses amplitude. These gains and losses combine the effects of optics, sensor noise, and vendor processing. Added high-frequency amplitude can come from noise or sharpening as well as from tissue structure. The curves therefore do not measure resolution. The same scanner also changed some tissues more than others. For example, AKOYA kept 45% of the high-frequency amplitude of AT2 in the cornea but only 17% in the aorta. A correction calibrated on one mix of tissues may therefore fit another mix poorly. Tissue changed the size of these effects but not the order of the most distinct scanners. Here, a scanner effect includes the whole acquisition pipeline (Supplementary Table S2). The scanner model name alone may not be enough to choose correction parameters.

Aligning images did not reliably align representations. Pix2Pix reproduced all measured image properties at three quarters of the locations, yet it moved UNI away from the target. Probes also still separated its outputs from real targets. Color–frequency correction improved image fidelity for both AKOYA and GT450 but moved UNI toward the target only for AKOYA. Matching color and frequency removed at most about a fifth of the PFM distance to the target. Most of the representation shift therefore remained after these global corrections. Whether a correction helped depended on the PFM, the tissue type, and whether a scanner had more or less high-frequency amplitude than the reference. Corruption benchmarks measure how far embeddings drift under synthetic changes (Coşar, 2026). Real paired targets add the direction of that drift. This direction decides whether a correction helps.

More complex correction was not consistently better. Reinhard with frequency matching was comparable to learned translation on several measures in PanNormal. A correction built from the measured scanner phenotypes is therefore worth testing alongside learned translation. Feature-level correction aligns paired embeddings directly, as in FEATMAP (Donle et al., 2026). It gave the largest gains within PanNormal and preserved tissue retrieval. However, it must be fitted for each PFM from paired data. Its ridge maps did not transfer to PLISM. After ComBat, a linear probe was close to chance, but a nonlinear probe still distinguished corrected from real target embeddings.

The controlled frequency changes also tracked robustness across models. Among eight models pretrained on WSI tiles, those whose representations moved least under a small frequency change were the most robust to real scanner differences. In general computer vision, robustness depends on which spatial frequencies a model relies on (Yin et al., 2019). Here, robustness tracked how far a model moved, not which band it preferred. UNI2-h and Virchow2 weighted opposite bands. The band ratio was unrelated to robustness. Models trained on natural images were less robust than their sensitivity predicted. They also had fewer neighbors that were the same tissue from another scanner. The sensitivity result links the wide differences in robustness reported for PFMs (de Jong et al., 2025; Kömen et al., 2026) to a response that can be measured with synthetic image changes. Virchow2, the least sensitive model, was trained on tiles from several magnifications (Zimmermann et al., 2024). Such training may teach a model to depend less on fine texture. Training choices that are often expected to improve robustness did not remove the scanner signal here. Spatial-transcriptomic fine-tuning (Hemker et al., 2026) increased the fraction of same-tissue, cross-scanner neighbors of UNI2-h. Its measured scanner detectability, distance, and frequency sensitivity changed little. In UNI2-h, the anchor thus seems to have made nearest neighbors more often the same tissue without reducing scanner sensitivity. In CONCH, the anchor did not change the robustness index. EXAONEPath uses stain normalization during pretraining and inference (Yun et al., 2024). However, it remained highly sensitive. One possible reason is that stain normalization does not address the frequency differences that scanners introduce. PLIP and CONCH, both trained with text, differed widely in robustness (robustness index 0.06 and 0.41). One difference is the starting point. The image encoder of CONCH was pretrained on WSI tiles before text alignment (Lu et al., 2024), whereas PLIP fine-tuned an encoder pretrained on natural images (Huang et al., 2023). That encoder was CLIP. CLIP already had most of PLIP’s same-scanner neighbors. In this pair of models, pathology image–text fine-tuning therefore changed this neighbor pattern only slightly.

Two features of pretraining suggest why scanner signals may persist. First, the scanners that produced the pretraining images can shape the frequency response. Changing only the share of images from two scanners shifted the band weighting of UNI and raised the band sensitivity of Virchow2. These changes were small and differed between models. Still, they suggest that pretraining data should be described by their scanners as well as by slides, tissue types, and institutions. Second, DINO and DINOv2 learn to make outputs agree across augmented views of the same image (Caron et al., 2021; Oquab et al., 2024). However, the evaluated augmentations did not reproduce real scanner differences, even when chosen with the real target. Blur also brought scanners together only by removing tissue structure. Real multi-scanner images of the same tissue, or augmentations built from measured scanner phenotypes, could narrow the gap between augmented and real scanner images. Frequency sensitivity could also be monitored during such training.

These results suggest five requirements for evaluating scanner correction. First, a real paired target is needed, because a smaller distance between scanners did not show that a correction reached the target. Second, image fidelity and representation alignment should be measured separately, because they did not move together. Pixel-wise measures such as SSIM were also far more sensitive to small misregistration than PFM distances (Supplementary Section B.3).

Third, whether corrected embeddings remain distinguishable from real targets should be tested with more than one kind of probe, because linear and nearest-neighbor probes disagreed after feature correction. Fourth, alignment should be checked in more than one PFM and in an external acquisition context, because effects reversed across both. Fifth, tissue information should be checked alongside alignment, because blur shrank the distance between scanners by erasing tissue structure.

This study has limitations. We used normal tissues, which differ from the tumor slides common in downstream applications. We also compared complete acquisition pipelines without separating scanner model, individual device, imaging time, and vendor processing. All models were evaluated as frozen encoders, and the augmentation analyses used UNI only. The relation between sensitivity and robustness rests on eight models pretrained on WSI tiles. These models also differ in architecture, data, and training objective. The relation is therefore an association that new models can test. It also did not extend to the robustness index of models trained on natural images. Frequency sensitivity was measured on images color-matched to each target scanner. Whether it can be measured from single-scanner images alone remains to be tested. The pretraining experiment shows that the scanners in pretraining data can change frequency weighting. It does not show how much of the differences between published models these scanners explain. Finally, coarse tissue classification was not affected by the raw scanner gap in our data (Supplementary Section B.7). The downstream benefit of correction should therefore be tested on tasks that depend on finer tissue features.

## 5. Conclusion

We imaged the same tissue on six scanners. This allowed us to decompose scanner batch effects into measurable color and spatial-frequency changes and to link these changes to PFM representations. Scanners changed color and spatial frequency by amounts that depended on tissue. Matching color and spatial frequency aligned images more reliably than representations. Probes distinguished corrected from real target embeddings after every image-level and feature-level correction. PFMs responded differently to frequency bands. Among models pretrained on WSI tiles, those that moved less under small frequency changes were more robust to real scanner differences. Scanner correction should therefore be judged against real paired targets on images, representations, and tissue information together.

## A. Supplementary Methods

### A.1. Dataset processing details

Supplementary Table S2. gives native resolution and acquisition magnification by scanner in both datasets. In local OD-gradient matching to other scanners, we excluded locations with a score below 0.5, a margin below 0.02 over an alternative peak, a best peak at the search boundary, or a difference of more than 2 pixels between the coordinate estimates of different matchers, as well as locations outside image boundaries. The 40 fitting locations per slide were the first four locations by source index in each of 10 spatial replicate groups. The 20 evaluation locations were the first two per group. PLISM scanners P, S210, and SQ are absent from PanNormal. PLISM does not include AKOYA. All PLISM images were acquired at 40×. In total, 18,739 PLISM locations were matched across all seven scanners. The evaluation set was drawn from these locations.

**Table S1.** PanNormal slides and fully matched locations per tissue type (37 tissue types, 103 slides, 95,780 locations). Every slide was imaged on all six scanners.

| Tissue type | Slides | Locations | Tissue type | Slides | Locations |
| --- | --- | --- | --- | --- | --- |
| Adrenal | 3 | 3,000 | Ovary | 2 | 2,000 |
| Aorta | 3 | 2,774 | Pancreas | 3 | 3,000 |
| Artery | 2 | 628 | Parathyroid | 3 | 3,000 |
| Bone marrow, growth plate | 3 | 3,000 | Pituitary | 3 | 2,847 |
| Brain, cerebellum | 3 | 3,000 | Prostate | 3 | 3,000 |
| Brain, cortex | 3 | 3,000 | Salivary gland | 3 | 3,000 |
| Breast | 2 | 1,980 | Skeletal muscle | 3 | 3,000 |
| Cervix, uterus | 2 | 2,000 | Skin | 3 | 2,145 |
| Choroid plexus | 2 | 2,000 | Small intestine | 4 | 4,000 |
| Esophagus | 3 | 2,344 | Smooth muscle | 1 | 890 |
| Eye (cornea) | 2 | 1,520 | Spinal cord | 3 | 3,000 |
| Fallopian tube | 3 | 2,828 | Spleen | 3 | 3,000 |
| Gall bladder | 3 | 1,819 | Stomach | 3 | 2,450 |
| Heart | 3 | 3,000 | Testis | 2 | 2,000 |
| Kidney (cortex/medulla) | 3 | 3,000 | Thymus | 3 | 3,000 |
| Large intestine (colon) | 3 | 3,000 | Thyroid gland | 3 | 3,000 |
| Liver | 3 | 3,000 | Tonsil | 2 | 2,000 |
| Lung | 3 | 2,131 | Urinary system | 5 | 4,424 |
| Lymph node | 2 | 2,000 |  |  |  |

**Table S2.** Native resolution and acquisition magnification of scanner models by manufacturer. All analysis images were converted to 0.5052 *μ*m/px.

| Scanner | Manufacturer | Model | Native MPP ($\mu\text{m}/\text{px}$) | Magnification |
| --- | --- | --- | --- | --- |
| **PanNormal** |  |  |  |  |
| AKOYA | Akoya Biosciences | PhenoCycler-Fusion | 0.4999 | 20× |
| S360 | Hamamatsu Photonics | NanoZoomer S360 | 0.4603 | 20× |
| S60 | Hamamatsu Photonics | NanoZoomer S60 | 0.4426 | 20× |
| AT2 | Leica Biosystems | Aperio AT2 | 0.5052 | 20× |
| GT450 | Leica Biosystems | Aperio GT 450 | 0.2624 | 40× |
| VERSA | Leica Biosystems | Aperio VERSA | 0.2742 | 20× |
| **PLISM** |  |  |  |  |
| S210 | Hamamatsu Photonics | NanoZoomer S210 | 0.2208 | 40× |
| S360 | Hamamatsu Photonics | NanoZoomer S360 | 0.2297 | 40× |
| S60 | Hamamatsu Photonics | NanoZoomer S60 | 0.2208 | 40× |
| SQ | Hamamatsu Photonics | NanoZoomer SQ | 0.2211 | 40× |
| AT2 | Leica Biosystems | Aperio AT2 | 0.2532 | 40× |
| GT450 | Leica Biosystems | Aperio GT 450 | 0.2621 | 40× |
| P | Philips | Ultra Fast Scanner | 0.2500 | 40× |

### A.2. Image properties and transfer curves

For the frequency properties, we averaged OD across RGB channels and removed the patch mean before computing the radial Fourier amplitude. CIELAB Δ*E*76 is the Euclidean distance between the two images’ mean *L*^∗^, *a*^∗^, and *b*^∗^ values,

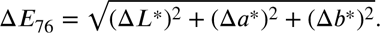

Zero means equal mean color. The measure does not assess edge or structural agreement. In PanNormal, slide-level contrasts were averaged within tissue type. In PLISM, core-level contrasts were averaged across stained sections. Transfer curves used the mean power spectrum over all matched locations of each slide in 72 radial bins. Each curve was expressed as the log_2_ amplitude ratio to AT2. Curves were averaged first within and then across tissue types. Coherence with AT2 was computed from the same paired spectra. Band averages of the curves reproduced the slide-level band summaries exactly.

### A.3. Scanner-by-tissue mixed-effects model

Let *desr* be the contrast of image measure *e* between target scanner *s* and the reference scanner on slide *r*, and let *t*(*r*) be the slide’s tissue type. For each measure, we fitted the linear mixed-effects model

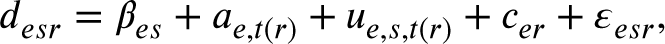

where *β_es_* is the scanner-specific mean difference (fixed), *a_e,t(r)_* the tissue effect shared across the five target scanners, *u_e,s,t(r)_* the scanner-by-tissue effect, *c_er_* the physical-slide effect, and *εesr* the residual. Each random effect was fitted by restricted maximum likelihood as an independent zero-mean component. The scanner-by-tissue term had a variance shared across the five target scanners. Variance fractions divided each variance by the sum of the variances of the shared tissue, scanner-by-tissue, slide, and residual effects. We tested tissue dependence against a reduced model with both *a* and *u* variances set to zero. We tested the scanner-by-tissue interaction against a reduced model with only the *u* variance set to zero. Both tests were restricted likelihood-ratio tests with 500 parametric bootstrap replicates. The smallest attainable *p* was therefore 0.002. The *p* values were adjusted by the Benjamini–Hochberg method across the 13 measures. The same model was applied to slide-level correction gains (Supplementary Section B.6).

### A.4. Image-level correction details

The 65 augmentation candidates consisted of the original image and 64 Sobol draws. The 257-candidate set added 192 draws over a wider range (Supplementary Table S3). For the oracle, we required source-gradient NCC of at least 0.90 (structure constraint). The upper limit on saturation fraction was 0.10 or the source value plus 0.05, whichever was greater. We also evaluated the 257-candidate set without the structure constraint. Candidates were selected at 40 locations per slide. SSIM, LPIPS–VGG16, and UNI distance were evaluated at the 20 evaluation locations. Augmentation improvement percentages were calculated relative to the original image at each location.

For single-factor trajectories, blur used *σ* = 0, 0.5, 1, 2, 3, 6 pixels. The unsharp mask added *α*(*I* − *Gσ*=1(*I*)) to image *I* for *α* = 0, 0.25, 0.5, 1, 2. Results were clipped to the RGB range. At each location, we subtracted the mean raw six-scanner UNI embedding. PCA was then fitted to 60 mean states (six scanners, two operations, five strengths each; blur strengths *σ* ≤ 3). The *σ* = 6 state was projected onto these axes. Cosine similarity between displacement directions was computed for 15 scanner pairs using, for each scanner, the mean UNI vector from the raw to the transformed image. The interscanner distance ratio was the mean UNI cosine distance between scanners at matched locations after transformation, divided by the same distance for raw images. We report the median over 15 pairs. Retained gradient energy was the grayscale-gradient energy relative to the original image. For tissue retrieval, slide-mean embeddings were queried against other physical slides. For the scanner probe, a class-balanced six-way linear probe used the locked folds.

Pix2Pix checkpoints were selected by paired-target pixel L1 on validation images. CycleGAN checkpoints were selected by comparing the distributions of the 10 image properties between generated images and the target domain. For the GT450 direction, we added an edge constraint to Pix2Pix: a weight of 100 on the mean L1 difference between horizontal and vertical grayscale gradients of source and generated images. Architecture, seed, 200-pass training, and checkpoint selection matched the original Pix2Pix. The two models were compared at 40 locations on held-out slides. Frequency matching applied scanner-specific radial gains to the Fourier amplitude of mean RGB OD. The gains were estimated from paired OD spectra on training slides and limited to 0.5–2.0. Phase and between-channel OD differences were preserved.

**Table S3.** Augmentation candidates. Each candidate drew every parameter uniformly from its range by a scrambled Sobol sequence. Additional Sobol dimensions decided whether the color, stain, blur, sharpening, and noise operations were applied. Blur and sharpening were mutually exclusive. The 65-candidate set consisted of the original image and 64 draws from the narrow ranges. The 257-candidate set added 192 draws from the wide ranges. Applied: share of candidates in which the parameter differed from the identity value.

| Parameter | Range |  | Applied (%) |  |
| --- | --- | --- | --- | --- |
|  | Narrow | Wide | 65 set | 257 set |
| Brightness factor | 0.80–1.20 | 0.40–1.60 | 83 | 84 |
| Contrast factor | 0.80–1.20 | 0.40–1.80 | 83 | 84 |
| Saturation factor | 0.80–1.20 | 0.40–1.80 | 83 | 84 |
| Gamma | 0.80–1.20 | 0.50–1.80 | 83 | 84 |
| H stain scale | 0.85–1.15 | 0.50–1.60 | 65 | 65 |
| E stain scale | 0.85–1.15 | 0.50–1.60 | 65 | 65 |
| Gaussian blur $\sigma$ (px) | 0.00–1.00 | 0.00–3.00 | 28 | 29 |
| Unsharp-mask amount $\alpha$ | 0.00–0.50 | 0.00–2.00 | 29 | 29 |
| Gaussian noise SD | 0.000–0.025 | 0.000–0.060 | 34 | 34 |

The locations in Fig. 3 were chosen by a fixed rule. For each direction, a location was eligible if three conditions held. First, for both image residual and UNI distance, every correction differed from raw in the same direction as the mean over the 2,060 evaluation locations. Second, color–frequency correction differed from Reinhard in the same direction as the mean. Third, the corrections with the lowest residual and the lowest distance were the same as in the mean (Pix2Pix and CycleGAN in both directions). There were 501 (AKOYA) and 41 (GT450) eligible locations. Among them, we show the one whose 10 values were closest to the means. Closeness was measured as the smallest sum of squared *z*-scores. The examples are therefore illustrative rather than typical of every location.

### A.5. Feature-level correction details

Feature correction used unit-normalized raw embeddings from 40 locations per training slide. Affine least squares (OLS) and ridge were fitted to matched source–target embeddings. The ridge penalty was chosen from 0.01, 0.1, and 1.0 times the mean diagonal of the training Gram matrix to minimize paired-target cosine distance in inner validation. ComBat used the target scanner as the reference batch, without tissue type in the fit. The fitted transformations were applied to the 20 locations of held-out slides and to PLISM without refitting.

### A.6. Model details

All models received the same 256 × 256 image covering 129.3 *μ*m. Images were resized to the 224 × 224 model input by bicubic interpolation for UNI, UNI2-h, and Virchow2. H-optimus-1 used bilinear interpolation. Band-wise OD changes were matched in 256 × 256 images before resizing. Frequency sensitivity therefore describes each encoder together with its released preprocessing. CONCH and SEAL-CONCH features were taken before the final projection. Projected features served as a secondary check. EXAONEPath was used as released: its Macenko normalizer with the bundled reference, resizing to 256, center cropping to 224, and ImageNet normalization. Macenko failed on 147 of 166,860 images. For these images, the released fallback (the input image) was kept. Excluding them did not change the results. The same model without Macenko normalization was evaluated as an off-label comparison. PLIP and DINOv2 used their released processors. We reproduced these processors on floating-point tensors so that small band changes were not lost to quantization. DINOv2’s processor center-crops 224 pixels of the 256-pixel field. CLIP ViT-B/32 was loaded and preprocessed exactly as PLIP, and its features were the projected image embeddings. ResNet50 (torchvision ImageNet weights) received the whole 256-pixel field resized to 224 pixels by antialiased bicubic interpolation and ImageNet normalization. We did not use the released resize-and-crop transform. This way, ResNet50 saw the same field as CLIP and PLIP. Its features were global-average-pooled layer-4 outputs. Features pooled after layer 3, as in CLAM (Lu et al., 2021), were analyzed as a secondary variant. CLIP and ResNet50 embedded re-rendered images. PLIP embeddings of the same renders agreed with the stored PLIP embeddings (cosine similarity ≥ 0.999 for every image). The center crops of EXAONEPath and DINOv2 reduce their effective field of view to 113.2 *μ*m.

### A.7. Detectability, robustness index, and tissue retrieval

Detectability used L2-normalized embeddings. In PanNormal, for each outer fold we used only the slides held out from that fold’s correction fit. We split these slides into five inner groups. We trained on four groups and tested on one.

Accuracy was averaged per location, then per slide with classes balanced within slide, then over 103 slides. In PLISM, corrections were fitted on PanNormal. We therefore used five folds of sections. The linear probe was a standardized logistic regression (*C* = 0.1, liblinear); the multilayer perceptron had one hidden layer of 256 units (ReLU, *α* = 10−3, early stopping on a 10% validation split, patience 10, at most 300 iterations); the *k*-nearest-neighbor probe used *k* = 15 and cosine distance. The best probe was the maximum of the three within each bootstrap resample. For the comparison of models (Supplementary Table S10), no correction was fitted. The same three probes separated raw reference-scanner from raw target-scanner embeddings, pooled over the five target scanners, in five folds of physical slides. These values therefore differ from the raw rows of Tables 1 and 2. Those rows used only slides held out from each correction fit.

For the PathoROB robustness index (Kömen et al., 2026), we pooled raw embeddings from all six scanners at the evaluation locations. Tissue type was the biological class. Scanner was the confounder. For each anchor, we found its *k* nearest neighbors, excluding neighbors from the same physical slide. We counted same-tissue neighbors from another scanner (SO) and other-tissue neighbors from the same scanner (OS). The index was ∑SO/∑(SO + OS). The sums ran over neighbor ranks 1, …, *k* and all anchors from the 36 tissue types with at least two slides. Following PathoROB, *k* was the value in {5, 10, 15, 20, 25, 30, 40, 50} that maximized *k*-nearest-neighbor balanced accuracy for tissue type. The chance level of 0.089 was computed exactly from the label counts. Tissue retrieval used slide-mean embeddings and a bank of real target-scanner slide means from other slides. In PLISM, core and section means were used with other sections as the bank. For the comparison of models, query and bank were raw reference-scanner slide means. For the blur trajectories, query and bank were slide means of the same scanner and transformation. These three variants give different raw values.

### A.8. Frequency-band manipulation and sensitivity

Band manipulation used the 20 evaluation locations per slide. At each location, we matched the color of the reference-scanner image to each of the five target scanners. We used Reinhard parameters from the training slides of the fold that contained the held-out slide. We converted this image to mean OD. We then increased or decreased the Fourier amplitude of one band at a time. This step used smooth tapering and reflection padding. It preserved phase and between-channel OD differences. For each color-corrected image, we took the smallest OD root-mean-square (RMS) of the three band components. We set the target change (dose) to 0.25 or 0.50 times that value. Band coefficients were adjusted until the OD RMS change of the final RGB image reached the target. The target direction was the sign of the frequency gain of each scanner, averaged over the band and estimated from training slides. Frequency sensitivity was the mean representation shift at dose 0.25 divided by the model’s mean between-tissue distance among raw AT2 embeddings. The band ratio was high-over low–mid-band sensitivity. Spearman correlations were computed on point estimates across the eight models. The partial correlation controlled for between-tissue distance. For each reference model, we computed the residual from the least-squares line across the eight models. Its CI came from refitting the line in every slide resample. The correlations across 12 models used 100,000 random permutations. All other permutation tests were exact.

### A.9. Continued pretraining on scanner mixtures

Continued pretraining used the locations of the four folds that were not held out (75,891 locations from 82 slides). The 21 slides of the remaining fold were used only for evaluation. Each location received a fixed uniform random number *u*. Let *p* be the AT2 share. A location contributed its AT2 image if *u* < *p* and its GT450 image otherwise. The assignments were therefore nested across shares. Every run saw the same tissue locations. We used the official DINOv2 implementation (DINO, iBOT, and KoLeo losses with Sinkhorn–Knopp centering) (Oquab et al., 2024). Its heads of 16,384 prototypes were newly initialized with the same seed in all runs. We kept its augmentations except Gaussian blur. We omitted blur so that the frequency differences between the two scanners reached the model. UNI was loaded into a ViT-L/16. Virchow2 was loaded into a ViT-H/14 with four register tokens. Both reproduced the released embeddings (cosine similarity above 0.999). Global crops were 224 pixels and local crops 96 (UNI) or 98 (Virchow2) pixels. The schedule was set for three epochs at batch size 256. The backbone was frozen for the first half epoch while the heads trained. The backbone learning rate then rose over a quarter epoch to 10−4 in the last block, with layer-wise decay 0.9. It then followed a cosine schedule. We analyzed the teacher backbone after the first epoch (296 iterations). Each share was run with two data orders. The two orders shared the image assignments. They differed in the order and augmentation of images.

Evaluation used the 20 evaluation locations of the 21 held-out slides, the same band manipulations as for the released models, and the released preprocessing of each model. Sensitivity was divided by the between-tissue distance of the same model. The band ratio therefore does not depend on the distance scale of a model. For each data order, we fitted a least-squares line of the band ratio on the AT2 share. The reported slope pools both data orders. CIs came from 2,000 resamples of the held-out slides, shared by all models. Tissue information was measured as nearest-neighbor tissue classification of held-out images from the four scanners absent from pretraining. The reference set was images of the training slides from the same scanner.

### A.10. Evaluation and uncertainty details

Let 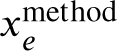 and 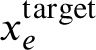 be the values of property *e* for a corrected image and its paired target. Let *σ*_e_ be the robust scale of property *e*. It is the interquartile range of the paired target–reference difference in *e*, divided by 1.349. It was computed over the fitting locations and five target scanners of the training slides. The standard deviation was used when this range was near zero. Image residual is

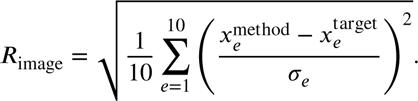

Coverage was the proportion of properties with 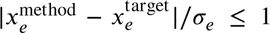. Joint coverage was the proportion of locations at which all 10 properties met this criterion. SSIM and LPIPS were computed on the 252 × 252 interior, excluding a 2-pixel boundary. Target-gradient NCC assessed agreement with target edges. Source-gradient NCC assessed preservation of original structure. Saturation fraction is the proportion of image values at the extremes of the representable range. Invented-edge fraction is the proportion of strong edges newly appearing where neither source nor target has an edge. For a raw value *R* and method value *M*, percentage improvement was 100(*M* −*R*)∕*R* for SSIM and 100(*R* − *M*)∕*R* for LPIPS and PFM distance. The between-tissue distance was the mean cosine distance over all location pairs from different tissue types. It was recomputed in every bootstrap resample. Retrieval CIs resampled query slides with the reference bank fixed.

## B. Supplementary Results

### B.1. Tissue-specific scanner effects (RQ1)

Supplementary Tables S4 and S5 give the scanner-specific mean differences and the variance components of the mixed-effects model for all 13 image measures. Tissue dependence and scanner-by-tissue interactions were detected for every measure (BH *q* ≤ 0.002). The scanner-by-tissue effect was largest for low–mid-frequency transfer (57% of the remaining variance) and smallest for gradient agreement with the reference image (9%). Supplementary Fig. S1 shows the tissue-specific scanner effect for four measures in every tissue type.

**Table S4.** Scanner-specific mean differences from the reference scanner (AT2) for all 13 PanNormal image measures, estimated with the linear mixed-effects model (Supplementary Methods). Tissue dependence and scanner × tissue interactions were significant for every measure (Benjamini–Hochberg *q* ≤ 0.002).

| Image measure | VERSA | AKOYA | GT450 | S360 | S60 |
| --- | --- | --- | --- | --- | --- |
| *Color and optical density* |  |  |  |  |  |
| $\Delta L^*$ | +7.436 | +6.187 | +17.943 | +1.703 | +8.819 |
| $\Delta a^*$ | −3.293 | +20.169 | −16.442 | +7.352 | −0.291 |
| $\Delta b^*$ | +1.372 | −17.160 | +5.658 | −6.565 | −0.336 |
| Mean OD ratio ($\log_2$) | −0.438 | −0.477 | −1.210 | −0.144 | −0.573 |
| *Contrast and edges* |  |  |  |  |  |
| $L^*$ contrast ratio ($\log_2$) | +0.043 | −0.732 | −0.634 | −0.135 | −0.267 |
| OD contrast ratio ($\log_2$) | −0.089 | −0.809 | −1.079 | −0.121 | −0.392 |
| Gradient energy ratio ($\log_2$) | −0.077 | −1.436 | −0.936 | −0.171 | −0.387 |
| *Spatial frequency* |  |  |  |  |  |
| Low–mid frequency transfer ($\log_2$) | +0.060 | −0.537 | +0.026 | −0.018 | +0.001 |
| Mid-frequency transfer ($\log_2$) | +0.005 | −1.354 | +0.242 | −0.114 | +0.008 |
| High-frequency transfer ($\log_2$) | −0.028 | −1.797 | +0.659 | −0.356 | −0.064 |
| *Supplementary measures* |  |  |  |  |  |
| 1–gradient NCC to AT2 | +0.658 | +0.371 | +0.328 | +0.263 | +0.303 |
| $\Delta$ tissue fraction | −0.048 | −0.001 | −0.055 | +0.014 | +0.015 |
| Laplacian variance ratio ($\log_2$) | +0.088 | −3.720 | −0.442 | −0.686 | −0.640 |

### B.2. Robustness of the high-frequency order to resampling (RQ1)

GT450 and VERSA images were downsampled about 1.9-fold and 1.8-fold to the common grid. In contrast, AKOYA images were barely resampled. To test whether the high-frequency order reflects resampling, we extracted native-resolution crops at the 20 evaluation locations. We resampled them to 0.5052 *μ*m/px with three methods: bicubic interpolation without anti-aliasing, anti-aliased Lanczos, and box integration over the AT2 pixel footprint. Under every method and in all 2,000 bootstrap resamples, GT450 had the highest high-frequency amplitude. AKOYA had the lowest. GT450’s log_2_ ratio stayed positive (+0.53 to +0.96, compared with +0.68 for the registration pipeline). VERSA and S60 changed sign with the method, and magnitudes shifted by −0.18 to +0.38. UNI distances to AT2 changed by at most 0.007 and kept their ranking.

### B.3. Registration floor and coherence (RQ1, RQ2)

To measure how much target distance remains when two images differ only by misregistration, we shifted AT2 images by 0.25 to 2 pixels. A 1-pixel shift is the size allowed by quality control. This shift produced 2–4% of the raw AT2–target distance in the four PFMs (UNI 2.8%, UNI2-h 3.5%, Virchow2 1.9%, H-optimus-1 3.6%). It also produced 3–6% of the best corrected distance. Pixel-wise similarity measures were far more sensitive. SSIM between an AT2 image and the same field shifted by 1 pixel was 0.729. With a 2-pixel shift, SSIM was 0.384. This value was below the raw AT2–target SSIM of 0.572. Coherence between paired images decayed toward zero at high frequency for every scanner. For VERSA, it had decayed toward zero by about 0.4 cycles/*μ*m (Supplementary Fig. S2). Paired high-frequency amplitude was thus not phase-aligned between scanners. This affects pixel-wise measures such as SSIM but not amplitude-based phenotypes.

**Table S5.** Variance components (%) of the linear mixed-effects model after removing scanner-specific mean differences, for all 13 PanNormal image measures. Shared tissue and shared slide are effects common to the five target scanners. Scanner × tissue is the tissue-specific scanner effect.

| Image measure | Shared tissue | Scanner × tissue | Shared slide | Residual |
| --- | --- | --- | --- | --- |
| *Color and optical density* |  |  |  |  |
| $\Delta L^*$ | 16.3 | 17.7 | 40.8 | 25.3 |
| $\Delta a^*$ | 0.0 | 29.9 | 0.0 | 70.1 |
| $\Delta b^*$ | 0.0 | 28.0 | 0.0 | 72.0 |
| Mean OD ratio ($\log_2$) | 0.0 | 32.9 | 1.8 | 65.3 |
| *Contrast and edges* |  |  |  |  |
| $L^*$ contrast ratio ($\log_2$) | 26.8 | 30.5 | 9.8 | 32.9 |
| OD contrast ratio ($\log_2$) | 0.0 | 43.2 | 0.0 | 56.8 |
| Gradient energy ratio ($\log_2$) | 2.6 | 35.9 | 0.9 | 60.6 |
| *Spatial frequency* |  |  |  |  |
| Low-mid frequency transfer ($\log_2$) | 0.0 | 57.4 | 0.0 | 42.6 |
| Mid-frequency transfer ($\log_2$) | 0.0 | 38.9 | 10.0 | 51.1 |
| High-frequency transfer ($\log_2$) | 0.0 | 20.1 | 21.3 | 58.6 |
| *Supplementary measures* |  |  |  |  |
| 1-gradient NCC to AT2 | 36.6 | 9.4 | 20.7 | 33.4 |
| $\Delta$ tissue fraction | 0.0 | 25.0 | 13.1 | 61.9 |
| Laplacian variance ratio ($\log_2$) | 11.6 | 25.7 | 25.6 | 37.0 |

### B.4. Augmentation and structure preservation (RQ2)

Supplementary Table S6. compares three augmentation oracles that selected candidates by location using real targets. In every scanner and all five folds, the global setting selected the original image. More candidates or removal of the structure constraint improved SSIM. However, UNI distance remained greater than raw. With 257 structure-preserving candidates, image residual fell from 1.313 for raw to 0.758. However, all 10 image properties were reproduced simultaneously at only 49.1% of locations. Balanced accuracy for distinguishing oracle images from real targets was 93.7%.

We examined 4,120 locations per scanner under the 257-candidate structure-preserving oracle. For 49.2% of AKOYA-directed locations, the selected combination was 1.07-fold brightness, 1.14-fold H stain, gamma 0.82, and blur *σ* = 0.65. For 35.4% of GT450-directed locations, the selected combination was 0.41-fold contrast, 0.74/0.67-fold H/E stain, and gamma 1.39. Frequently selected combinations thus differed by scanner.

On the 20 evaluation locations, blur and sharpening displacement directions had median cosine similarities of 0.772 (blur *σ* = 3) and 0.697 (sharpening *α* = 2) across scanner pairs. At *σ* = 6, the median interscanner distance ratio was 0.551. The median retained gradient energy was 0.98%. Same-tissue nearest-neighbor retrieval fell from 64.2% to 45.1%. In contrast, the six-way scanner probe fell only from 0.994 to 0.967. Symmetric low-pass filtering of all six scanner images showed the same pattern. The strongest filter reduced UNI AT2–target distance from 0.214 to 0.139. Meanwhile, a tissue probe lost 6 percentage points of balanced accuracy. A scanner probe stayed at 0.997.

**Table S6.** Percentage changes from raw for PanNormal augmentations selected by location using real paired images. Positive values indicate improvement. Values are means across 103 slides.

| Candidate selection | SSIM improvement (%) ↑ | LPIPS reduction (%) ↑ | UNI distance reduction (%) ↑ |
| --- | --- | --- | --- |
| 65, structure constraint | +8.1 | +7.7 | −1.1 |
| 257, structure constraint | +9.1 | +7.0 | −3.9 |
| 257, unconstrained | +11.3 | +8.6 | −2.2 |

**Figure S1:**
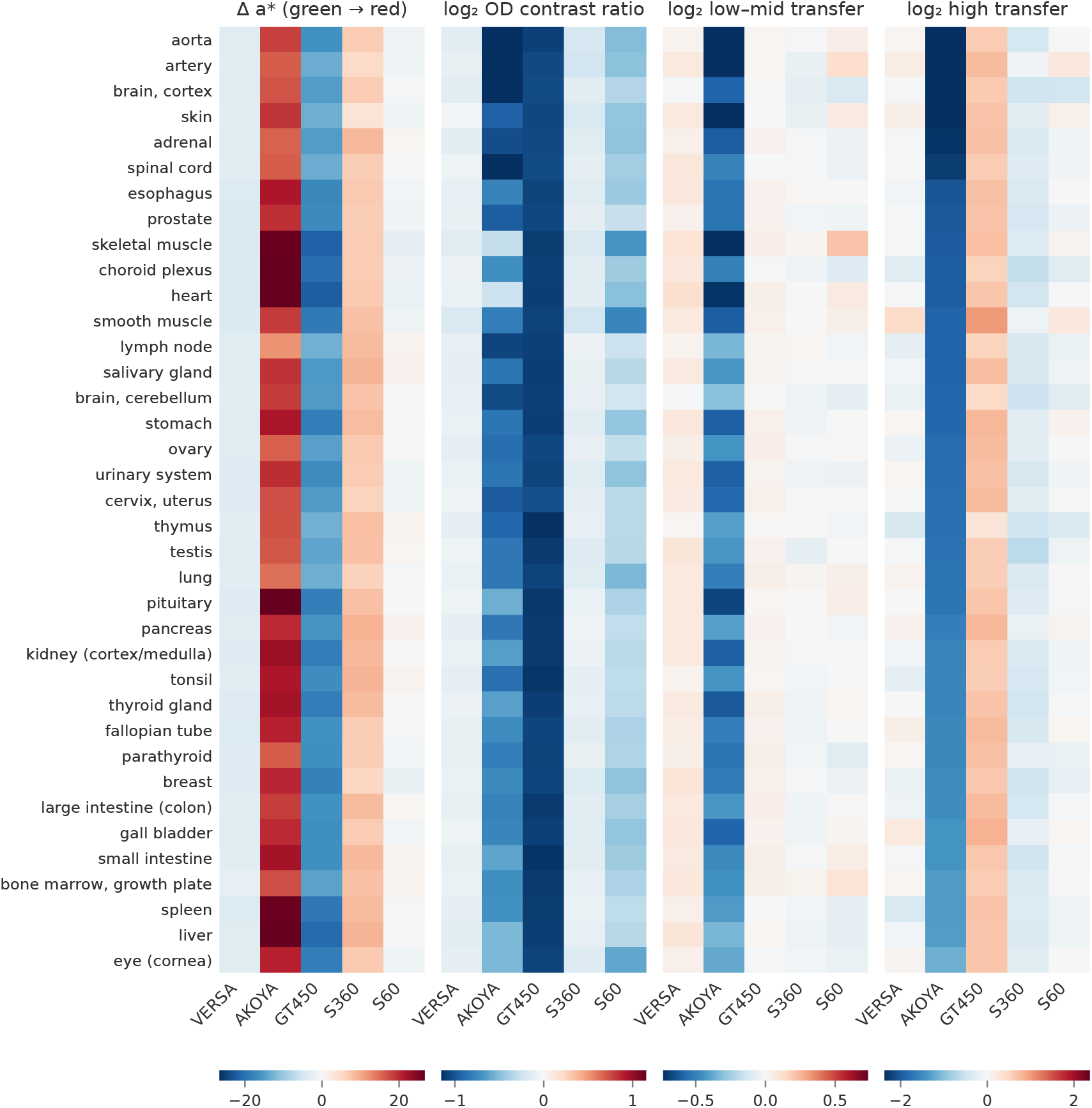
Tissue-specific scanner effects relative to AT2 in PanNormal. Panels show four measures: Δ*a*^∗^, log_2_ OD contrast ratio, and log_2_ low–mid and high-frequency transfer. Each cell is the mean contrast of one scanner over the slides of one tissue type. Rows are ordered by AKOYA high-frequency transfer. Color scales are centered at zero.

### B.5. Image-level correction details (RQ2)

Supplementary Table S7. gives image similarity and UNI distance after color-only, frequency-only, and combined correction in both datasets. Frequency matching alone improved all three PanNormal measures less than color correction alone. In PLISM, it worsened all three. In PanNormal, adding frequency matching reduced image residual from 0.698 to 0.499 (paired reduction 0.199, 95% CI 0.191–0.207). In contrast, the mean additional UNI gain was 0.0004 (95% CI −0.0009 to 0.0016). The additional gain was +0.0167 (95% CI 0.0105–0.0226) for AKOYA and −0.0153 (−0.0180 to −0.0127) for GT450, and −0.0070 (−0.0080 to −0.0060) across the three PLISM directions. For AKOYA, adding frequency matching after Reinhard raised target-gradient NCC from 0.579 to 0.633, lowered source-gradient NCC from 0.851 to 0.812, and changed same-tissue retrieval by +0.7 percentage points (95% CI −3.0 to 4.9).

Pix2Pix reduced image residual from 1.313 to 0.404 and raised joint coverage from 9.2% to 75.4%. CycleGAN reduced image residual from 1.313 to 0.550 and raised joint coverage from 9.2% to 60.5%. However, Pix2Pix kept less of the source structure (source-gradient NCC 0.644, vs 0.894 for CycleGAN). It also did not move closer to the target edges (target-gradient NCC 0.611, vs 0.620 for raw). Adding an edge constraint to GT450 Pix2Pix raised SSIM from 0.693 to 0.711, source-gradient NCC from 0.622 to 0.650, and target-gradient NCC from 0.457 to 0.478, and reduced invented-edge fraction from 0.0056 to 0.0043. However, UNI target gain did not improve (paired difference −0.007, 95% CI −0.017 to 0.004).

**Figure S2:**
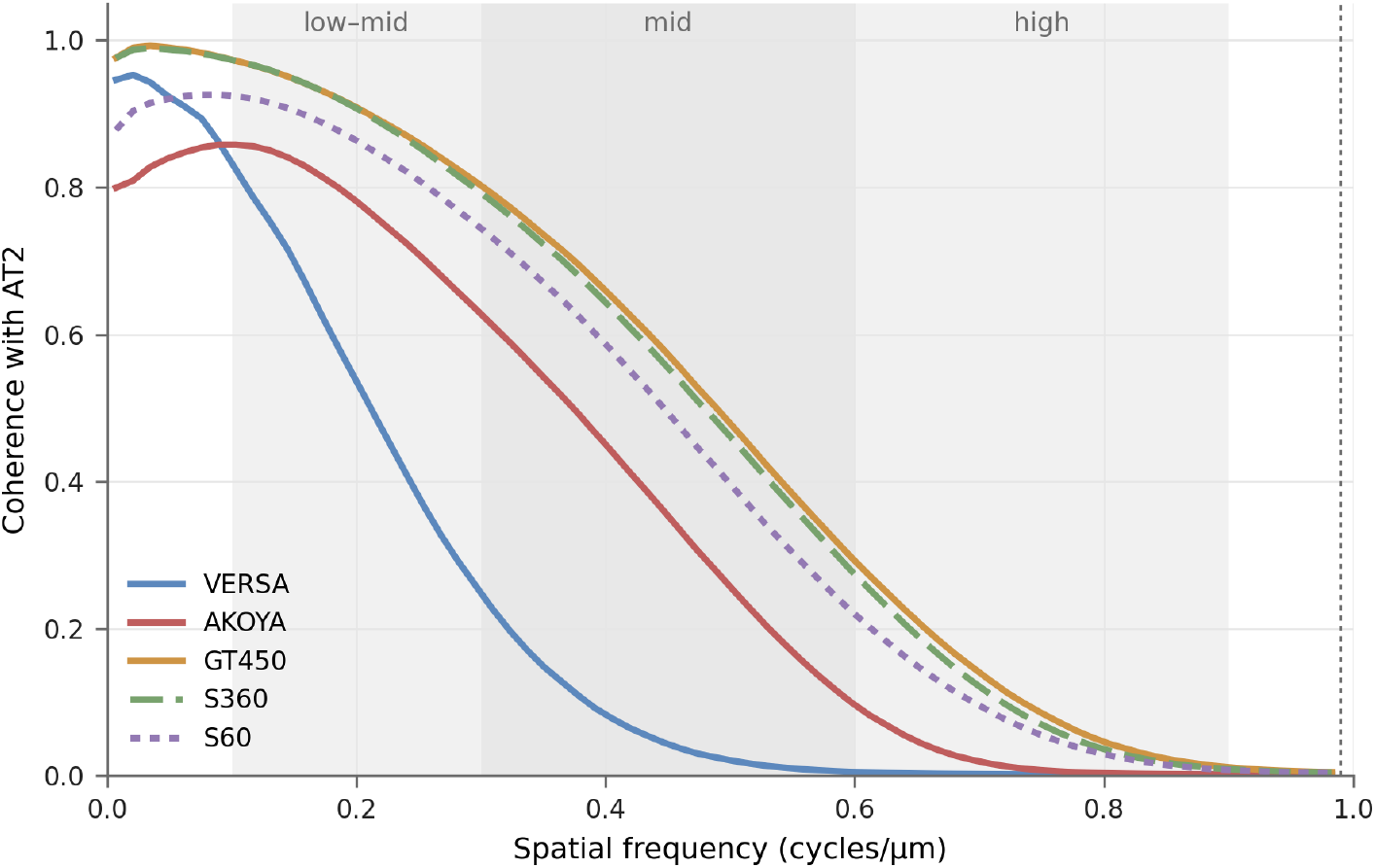
Coherence between paired target-scanner and AT2 images across spatial frequency, averaged over tissue types. Shaded regions mark the three analysis bands. The dotted line marks the AT2 Nyquist frequency.

**Figure S3:**
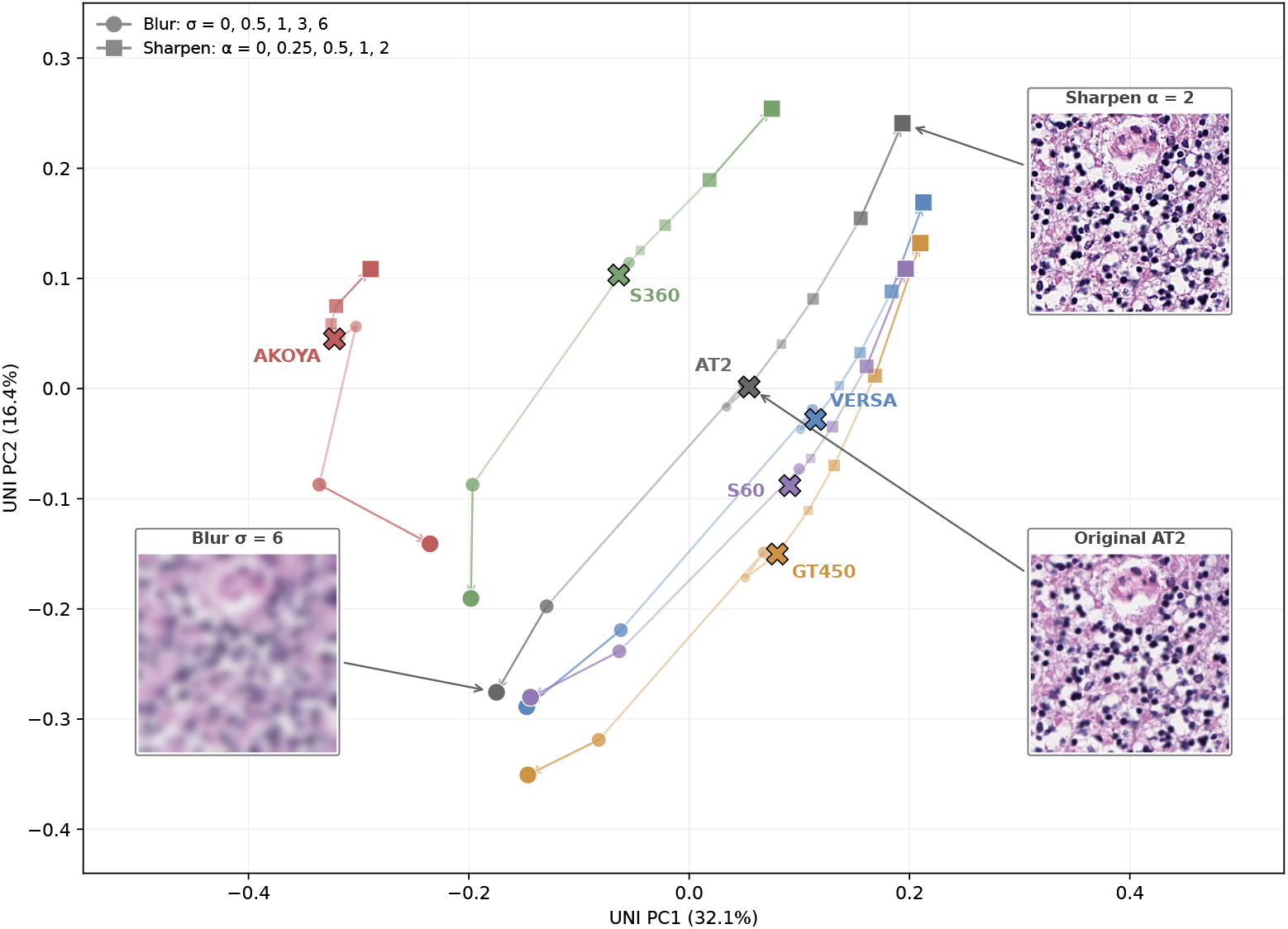
Blur and sharpening move UNI representations of all six scanners along shared directions. The plot shows PCA of centered embeddings. Color identifies scanner. Circles indicate blur, squares indicate sharpening, and X marks the original image. Marker size indicates strength (*σ* = 2 was included in the PCA fit but is not shown). Insets show the original image, blur (*σ* = 6), and sharpening (*α* = 2) at the same AT2 location. Twenty matched locations were used on each of 103 slides.

**Table S7.** Image similarity and UNI distance after color and frequency correction in PanNormal and PLISM (mean±SD). Color denotes Reinhard. Frequency denotes frequency matching. Δ Raw is the percentage improvement from raw. Positive values indicate improvement.

| Correction | SSIM ↑ | | LPIPS-VGG16 ↓ | | UNI distance ↓ | |
| --- | --- | --- | --- | --- | --- | --- |
| | Mean ± SD | $\Delta$ Raw (%) ↑ | Mean ± SD | $\Delta$ Raw (%) ↑ | Mean ± SD | $\Delta$ Raw (%) ↑ |
| **PanNormal** |  |  |  |  |  |  |
| Raw | 0.572 ± 0.074 | Ref. | 0.232 ± 0.026 | Ref. | 0.2136 ± 0.0365 | Ref. |
| Color | 0.620 ± 0.071 | +8.4 | 0.202 ± 0.025 | +12.8 | 0.1798 ± 0.0334 | +15.9 |
| Frequency | 0.597 ± 0.071 | +4.2 | 0.218 ± 0.026 | +5.9 | 0.2049 ± 0.0343 | +4.1 |
| Color + frequency | 0.626 ± 0.069 | +9.5 | 0.192 ± 0.024 | +17.2 | 0.1794 ± 0.0316 | +16.0 |
| **PLISM** |  |  |  |  |  |  |
| Raw | 0.781 ± 0.042 | Ref. | 0.161 ± 0.021 | Ref. | 0.1713 ± 0.0215 | Ref. |
| Color | 0.817 ± 0.039 | +4.6 | 0.153 ± 0.023 | +4.8 | 0.1754 ± 0.0233 | -2.4 |
| Frequency | 0.766 ± 0.041 | -1.9 | 0.164 ± 0.020 | -2.1 | 0.1895 ± 0.0219 | -10.6 |
| Color + frequency | 0.809 ± 0.039 | +3.6 | 0.156 ± 0.022 | +3.1 | 0.1824 ± 0.0229 | -6.5 |

### B.6. Tissue dependence of correction gains (RQ2)

We applied the scanner-by-tissue mixed-effects model to the slide-level target gains of Reinhard, color–frequency correction, the frequency increment over Reinhard, ridge affine, and ComBat in all four PFMs. We used Benjamini– Hochberg adjustment across the five methods within each PFM. Gains depended on tissue type in all 20 method–PFM combinations (BH *q* ≤ 0.006). The scanner-by-tissue interaction was detected in all 20 (*q* ≤ 0.034). The scanner-by-tissue effect explained 8–43% of the remaining variance. For AKOYA, the frequency increment was larger in tissues with a stronger high-frequency deficit: Spearman *ρ* = −0.57 in UNI, −0.35 in Virchow2 and H-optimus-1, and −0.16 in UNI2-h. No association was found for GT450.

### B.7. Downstream tissue classification (RQ2)

For each PFM and target scanner, we built tissue-type centroids from real target-scanner embeddings of all other slides. We then classified the held-out slide’s locations under the real target, raw AT2, and every corrected condition. Raw AT2 images were classified as well as the real targets. In none of the 20 PanNormal or 12 PLISM combinations of PFM and scanner did the accuracy gap between raw AT2 and the real target reach 2 percentage points (median gap −0.9 and +0.2 points). Several corrections lowered tissue accuracy instead. Pix2Pix lowered the tissue accuracy of Virchow2 from 52.0% to 43.8%. Macenko lowered that of UNI from 57.5% to 51.9%. In PLISM, Vahadane, Pix2Pix, and CycleGAN lowered the tissue accuracy of Virchow2 by 8–10 points.

### B.8. Detectability by probe and feature correction scope (RQ2)

Supplementary Table S8. gives detectability under each probe. Without the within-fold design, the linear probe fell below chance after ComBat in all four PFMs (0.29–0.36) and after affine OLS in UNI2-h and H-optimus-1 (0.35 and 0.41). The reason was that correction maps for one fold were fitted on targets that served as the classifier’s test fold. Within-fold values were 0.55–0.60 for ComBat and 0.58–0.74 for OLS. After ComBat and OLS, the multilayer perceptron detected the target scanner better than the linear probe in all four PFMs (differences +0.04 to +0.07 and +0.02 to +0.03). The *k*-nearest-neighbor probe detected ComBat-corrected embeddings at 0.62–0.80. After ridge, the linear probe detected the scanner at 0.67–0.87. In contrast, the *k*-nearest-neighbor probe was at 0.54–0.58. In PLISM, every method stayed at 0.99 or higher under the linear probe.

Supplementary Table S9. gives the target distances behind Table 2. In PanNormal, UNI ridge affine reduced raw distance from 0.2136 to 0.1403 (34.3%). In PLISM, it increased raw distance from 0.1713 to 0.2333 (36.2%). The paired difference in PanNormal target gain between ridge affine and color–frequency correction was +0.0390 (95% CI 0.0330–0.0448). For AKOYA, ridge affine and color–frequency correction had similar gains of +0.1557 and +0.1493.

**Table S8.**
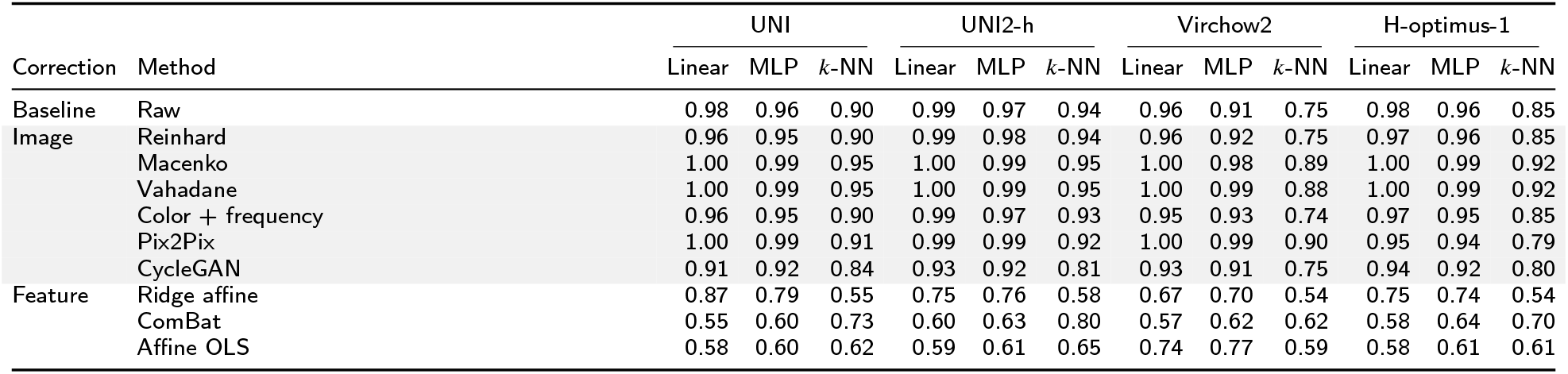
How well each probe distinguishes corrected from real target embeddings in PanNormal. Balanced accuracy of linear, MLP, and *k*-NN probes separating corrected embeddings from real target embeddings (0.5 = chance). Probes were trained and tested within the slides held out from each correction fit. Shaded rows are image-level corrections. In PLISM, every method stayed at 0.99 or higher under the linear probe.

Ridge affine worsened the VERSA and S360 directions for Virchow2 and the S360 direction for H-optimus-1. In PLISM, ComBat gain was positive for all four PFMs in the GT450 direction. It was negative for all four in the S360 direction. Affine OLS fitted on 20 instead of 40 locations per slide had a mean PanNormal gain of −0.0461 instead of +0.0552. Tissue retrieval after ridge correction ranged from 62.3% to 66.8% across PFMs. This was similar to the real-target range of 61.4%–67.4%.

Correction in the reverse direction, from each target scanner to AT2, showed the same pattern. Ridge affine, OLS, and Procrustes maps had positive mean gains within PanNormal (+0.071, +0.055, and +0.056). They had negative gains in PLISM (−0.053, −0.083, and −0.050). For GT450, we evaluated 40 locations per slide. In the reverse direction, Pix2Pix and CycleGAN moved UNI away from the target in both datasets. They moved UNI further away than in the forward direction.

**Table S9.**
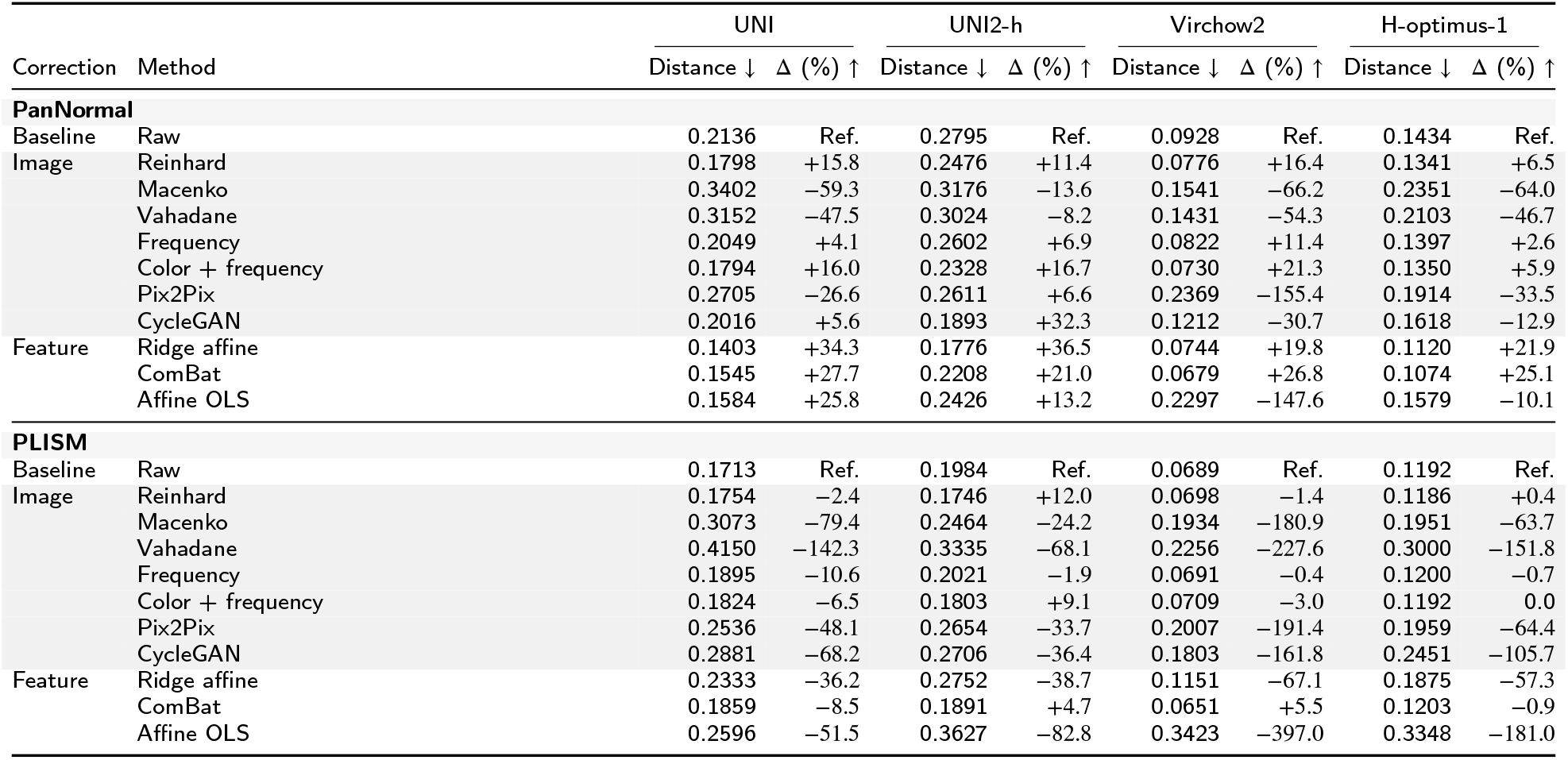
PFM cosine distance to the real paired target after each correction, and its change from raw (Δ, %; positive = closer). PanNormal: 103 slides, five directions; PLISM: 13 sections, three directions, with corrections fitted in PanNormal and applied without refitting. Shaded rows are image-level corrections. Frequency: frequency matching alone.

| Correction | Method | UNI |  | UNI2-h |  | Virchow2 |  | H-optimus-1 |  |
| --- | --- | --- | --- | --- | --- | --- | --- | --- | --- |
| | | Distance ↓ | $\Delta$ (%) ↑ | Distance ↓ | $\Delta$ (%) ↑ | Distance ↓ | $\Delta$ (%) ↑ | Distance ↓ | $\Delta$ (%) ↑ |
| PanNormal |  |  |  |  |  |  |  |  |  |
| Baseline | Raw | 0.2136 | Ref. | 0.2795 | Ref. | 0.0928 | Ref. | 0.1434 | Ref. |
| Image | Reinhard | 0.1798 | +15.8 | 0.2476 | +11.4 | 0.0776 | +16.4 | 0.1341 | +6.5 |
|  | Macenko | 0.3402 | -59.3 | 0.3176 | -13.6 | 0.1541 | -66.2 | 0.2351 | -64.0 |
|  | Vahadane | 0.3152 | -47.5 | 0.3024 | -8.2 | 0.1431 | -54.3 | 0.2103 | -46.7 |
|  | Frequency | 0.2049 | +4.1 | 0.2602 | +6.9 | 0.0822 | +11.4 | 0.1397 | +2.6 |
|  | Color + frequency | 0.1794 | +16.0 | 0.2328 | +16.7 | 0.0730 | +21.3 | 0.1350 | +5.9 |
|  | Pix2Pix | 0.2705 | -26.6 | 0.2611 | +6.6 | 0.2369 | -155.4 | 0.1914 | -33.5 |
|  | CycleGAN | 0.2016 | +5.6 | 0.1893 | +32.3 | 0.1212 | -30.7 | 0.1618 | -12.9 |
| Feature | Ridge affine | 0.1403 | +34.3 | 0.1776 | +36.5 | 0.0744 | +19.8 | 0.1120 | +21.9 |
|  | ComBat | 0.1545 | +27.7 | 0.2208 | +21.0 | 0.0679 | +26.8 | 0.1074 | +25.1 |
|  | Affine OLS | 0.1584 | +25.8 | 0.2426 | +13.2 | 0.2297 | -147.6 | 0.1579 | -10.1 |
| PLISM |  |  |  |  |  |  |  |  |  |
| Baseline | Raw | 0.1713 | Ref. | 0.1984 | Ref. | 0.0689 | Ref. | 0.1192 | Ref. |
| Image | Reinhard | 0.1754 | -2.4 | 0.1746 | +12.0 | 0.0698 | -1.4 | 0.1186 | +0.4 |
|  | Macenko | 0.3073 | -79.4 | 0.2464 | -24.2 | 0.1934 | -180.9 | 0.1951 | -63.7 |
|  | Vahadane | 0.4150 | -142.3 | 0.3335 | -68.1 | 0.2256 | -227.6 | 0.3000 | -151.8 |
|  | Frequency | 0.1895 | -10.6 | 0.2021 | -1.9 | 0.0691 | -0.4 | 0.1200 | -0.7 |
|  | Color + frequency | 0.1824 | -6.5 | 0.1803 | +9.1 | 0.0709 | -3.0 | 0.1192 | 0.0 |
|  | Pix2Pix | 0.2536 | -48.1 | 0.2654 | -33.7 | 0.2007 | -191.4 | 0.1959 | -64.4 |
|  | CycleGAN | 0.2881 | -68.2 | 0.2706 | -36.4 | 0.1803 | -161.8 | 0.2451 | -105.7 |
| Feature | Ridge affine | 0.2333 | -36.2 | 0.2752 | -38.7 | 0.1151 | -67.1 | 0.1875 | -57.3 |
|  | ComBat | 0.1859 | -8.5 | 0.1891 | +4.7 | 0.0651 | +5.5 | 0.1203 | -0.9 |
|  | Affine OLS | 0.2596 | -51.5 | 0.3627 | -82.8 | 0.3423 | -397.0 | 0.3348 | -181.0 |

### B.9. Frequency-band manipulation in four PFMs (RQ3)

At dose 0.25, the UNI shift was 0.0056 for the high band and 0.0044 and 0.0043 for the low–mid and mid bands. At dose 0.50, it was 0.0210, 0.0173, and 0.0168. High-band manipulation in the target direction gave a mean UNI gain of −0.0009 (95% CI −0.0012 to −0.0006) at dose 0.25 and −0.0086 (−0.0096 to −0.0076) at dose 0.50. A UNI representation shift alone did not indicate movement toward the real paired target.

The same manipulations were applied to UNI2-h, Virchow2, and H-optimus-1 (Fig. 5; Supplementary Fig. S4). At dose 0.25, the difference in representation shift between the high and low–mid bands (high minus low–mid) was +0.0012 for UNI (95% CI +0.0008 to +0.0016), +0.0031 for UNI2-h (+0.0025 to +0.0038), −0.0006 for Virchow2 (−0.0007 to −0.0004), and −0.0019 for H-optimus-1 (−0.0023 to −0.0015). The high-band gain in the target direction was positive for UNI2-h (+0.0053, 95% CI +0.0044 to +0.0062), close to zero for Virchow2 (+0.0001), and negative for H-optimus-1 (−0.0010).

**Figure S4:**
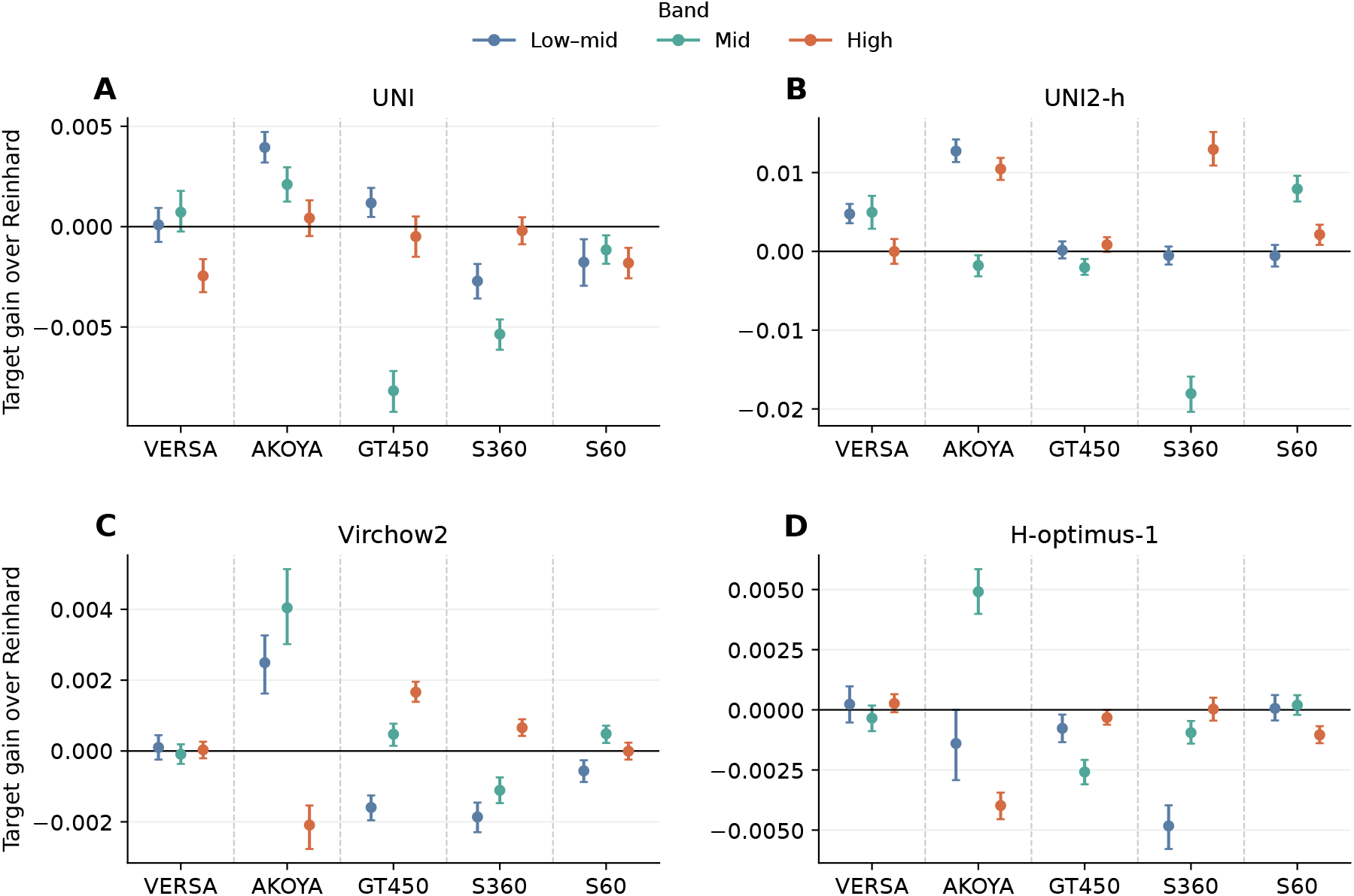
Additional gain toward the real paired target, relative to Reinhard alone, for **(A)** UNI, **(B)** UNI2-h, **(C)** Virchow2, and **(D)** H-optimus-1. One frequency band was changed in the direction of the target scanner (dose 0.25). Positive values indicate approach to the target. Dashed lines separate target scanners. Points and error bars show means and bootstrap 95% CIs across 103 PanNormal slides. Each slide contributed 20 matched locations.

### B.10. Training signal, data source, and scanner robustness (RQ3)

Supplementary Table S10. gives sensitivity, robustness, and tissue information for all model variants. We tested four explanations of scanner sensitivity. First, a non-image anchor did not reduce scanner sensitivity. SEAL changed normalized scanner distance by −0.013 for UNI2-h and +0.026 for CONCH. It left detectability and high-band sensitivity unchanged. However, SEAL did raise the robustness index of UNI2-h by 0.092 (95% CI 0.078–0.104). UNI2-h then had more same-tissue neighbors from other scanners (0.205 vs 0.151). SEAL left the robustness index of CONCH unchanged (−0.002). Second, we compared models whose final training used images other than WSI tiles (CONCH, PLIP, and DINOv2) with the seven other models. They were not lower on normalized scanner distance (difference +0.005). They had a lower robustness index (−0.13) and tissue retrieval (−0.19). The image encoder of CONCH was pretrained on WSI tiles before text alignment. CONCH therefore also enters the eight models pretrained on WSI tiles of the correlation analysis. Third, image-only models pretrained on WSI tiles had a slightly larger distance reduction from frequency matching and a slightly larger high-band sensitivity than the other models. Fourth, EXAONEPath as released had the largest normalized scanner distance (0.476; 0.378 without Macenko normalization), a larger distance reduction from Reinhard than the other models pretrained on WSI tiles (0.246 vs 0.125), and a low robustness index (0.18). Without Macenko normalization, its robustness index was lower (0.11). However, this unnormalized input differs from the normalized tiles on which the model was pretrained.

Across the eight models pretrained on WSI tiles, the relation between sensitivity and robustness held for every band. Spearman *ρ* with the robustness index was −0.76 (95% CI −0.79 to −0.57; permutation *p* = 0.037) for the low–mid band, −0.88 (−0.90 to −0.62; *p* = 0.007) for the mid band, and −0.90 (−0.98 to −0.74; *p* = 0.005) for the high band, and −0.95 (*p* = 0.001) for the high band at dose 0.50. The partial correlation controlling for between-tissue distance was −0.90. Two pairs of the eight models share a backbone (UNI2-h and SEAL-UNI2; CONCH and SEAL-CONCH). We then kept one model of each pair. With UNI2-h kept, the high-band *ρ* with the robustness index was −1.00 (exact permutation *p* = 0.003). With SEAL-UNI2 kept, it was −0.94 (*p* = 0.017). These values did not depend on which CONCH variant was kept. With PLIP and DINOv2 added to the eight models, the high-band *ρ* with the robustness index was −0.72 (*p* = 0.023). With CLIP and ResNet50 also added, it was −0.51 (*p* = 0.09). Across these 12 models, high-band sensitivity was related to normalized scanner distance with *ρ* = +0.83 (*p* = 0.001). The band ratio was not related to the robustness index (*ρ* = −0.19, *p* = 0.66) or to normalized scanner distance (*ρ* = +0.10). Best-probe detectability was at its ceiling and not related to sensitivity.

Supplementary Fig. S5 separates tissue information from scanner distance. PLIP had low tissue retrieval and a large normalized scanner distance. Of its nearest neighbors, 63% were other tissues from the same scanner, and only 4% were the same tissue from another scanner. CONCH, also trained with text, had 30% and 21%, respectively. It ranked among the more robust models. The low robustness index of PLIP therefore reflects a representation organized by scanner rather than by tissue.

PLIP was fine-tuned from CLIP ViT-B/32. Comparing the two models showed how much of this organization was present before fine-tuning. CLIP had a robustness index of 0.09 (PLIP 0.06; PLIP − CLIP difference −0.03, 95% CI −0.05 to −0.01). It had 58% other-tissue, same-scanner neighbors (PLIP 63%). On these two measures, 91% and 86% of the gap between PLIP and the median model pretrained on WSI tiles was already present in CLIP. Fine-tuning did increase normalized scanner distance, from 0.36 in CLIP to 0.46 in PLIP (difference +0.10, 95% CI +0.08 to +0.13). The reason was that the distance between scanners grew more than the distance between tissues. CLIP was about 1.8 times as sensitive as PLIP in the low–mid and mid bands. In the high band, the two models were similar. ResNet50 had a robustness index of 0.17, the lowest detectability of all models (0.85), and the lowest tissue retrieval (23%). However, its fraction of other-tissue, same-scanner neighbors (45%) lay within the range of the models pretrained on WSI tiles. Its low index therefore reflects weak tissue information rather than scanner organization. Features pooled after layer 3, as in CLAM, had a lower index (0.12) and more same-scanner neighbors (54%). Every reference model lay below the least-squares line of the robustness index on high-band sensitivity across the eight models pretrained on WSI tiles (residuals −0.14 to −0.33, all outside the range of the eight). On normalized scanner distance, DINOv2 and ResNet50 lay within that range. In contrast, CLIP (+0.13) and PLIP (+0.22) lay above it.

### B.11. Scanners in the pretraining data (RQ3)

In UNI, low–mid sensitivity rose with the AT2 share (slope +0.0003, 95% CI +0.0002 to +0.0004; *ρ* = +0.92). High-band sensitivity did not (−0.0000, −0.0002 to +0.0001; *ρ* = −0.18). In the two data orders, the band ratio was 1.38 and 1.37 with GT450 images only and 1.28 and 1.29 with AT2 images only. In each order, it decreased monotonically with the AT2 share (Spearman *ρ* = −1.00). In Virchow2, both low–mid (+0.0005, +0.0004 to +0.0006) and high-band sensitivity (+0.0004, +0.0003 to +0.0004) rose with the AT2 share. The band ratio stayed between 0.68 and 0.73 (Supplementary Fig. S6). The slope of the band ratio on the AT2 share, pooled over both data orders, was −0.093 (95% CI −0.116 to −0.075) in UNI and +0.006 (−0.004 to +0.015) in Virchow2. Normalized scanner distance from AT2 to the five other scanners rose with the AT2 share in both models (slope +0.011, 95% CI +0.009 to +0.014, in UNI and +0.010, +0.008 to +0.012, in Virchow2).

Continued pretraining also changed both models regardless of the share. Normalized scanner distance rose from 0.247 to 0.261–0.273 in UNI and from 0.128 to 0.179–0.192 in Virchow2. Nearest-neighbor tissue classification on the four scanners absent from pretraining fell from 0.569 to 0.540–0.556 in UNI and from 0.553 to 0.483–0.510 in Virchow2. Tissue classification also fell slightly with the AT2 share (slope −0.011 in UNI and −0.017 in Virchow2). We therefore compared runs with each other rather than with the released models.

**Table S10.** Frequency sensitivity, scanner robustness, and tissue information for all model variants (PanNormal, raw images). Sensitivity: representation shift under an equal high-band change, divided by the between-tissue distance. Scanner robustness: normalized scanner distance, best-probe detectability separating raw reference-scanner from raw target-scanner embeddings without correction (Det.), and PathoROB robustness index (RI; chance 0.089 in this dataset). Tissue: same-tissue retrieval among raw reference-scanner images of other slides (macro recall). This retrieval therefore differs from the target-scanner retrieval of Table 1. Distance reduction: fraction of the raw target distance removed by Reinhard, and the further fraction removed by adding frequency matching. The eight models pretrained on WSI tiles are listed first, followed by the reference models. Each group is ordered by RI. CLIP ViT-B/32 is the natural-image model from which PLIP was fine-tuned. ResNet50 features are the pooled layer-4 features of the ImageNet-supervised network.

| Model | Training signal and data | Sensitivity | Scanner robustness |  |  |  | Tissue | Distance reduction |  |
| --- | --- | --- | --- | --- | --- | --- | --- | --- | --- |
|  |  | High band ↓ | Distance ↓ | Det. ↓ | RI ↑ |  | Retrieval (%) ↑ | Reinhard | +Frequency |
| Virchow2 | Image-only, WSI | 0.0014 | 0.122 | 0.996 | 0.53 |  | 64.0 | 0.16 | 0.049 |
| H-optimus-1 | Image-only, WSI | 0.0019 | 0.160 | 0.997 | 0.51 |  | 66.9 | 0.06 | -0.006 |
| CONCH | Text anchor (figures) | 0.0040 | 0.139 | 0.973 | 0.41 |  | 58.7 | 0.21 | 0.018 |
| SEAL-CONCH | ST anchor | 0.0037 | 0.166 | 0.978 | 0.41 |  | 60.3 | 0.27 | 0.050 |
| SEAL-UNI2 | ST anchor | 0.0083 | 0.300 | 0.996 | 0.39 |  | 66.4 | 0.12 | 0.062 |
| UNI | Image-only, WSI | 0.0064 | 0.243 | 0.992 | 0.34 |  | 61.5 | 0.16 | 0.002 |
| UNI2-h | Image-only, WSI | 0.0083 | 0.313 | 0.995 | 0.30 |  | 65.5 | 0.11 | 0.053 |
| EXAONEPath | Image-only, WSI + stain norm. | 0.0100 | 0.476 | 0.991 | 0.18 |  | 50.6 | 0.25 | 0.036 |
| DINOv2 | Image-only, natural images | 0.0046 | 0.178 | 0.957 | 0.27 |  | 36.3 | 0.17 | 0.002 |
| ResNet50 | Supervised, ImageNet | 0.0039 | 0.234 | 0.847 | 0.17 |  | 23.2 | 0.41 | 0.050 |
| EXAONEPath (no Macenko) | Same, without Macenko | 0.0085 | 0.378 | 0.995 | 0.11 |  | 47.9 | 0.19 | 0.019 |
| CLIP ViT-B/32 | Text anchor (web), natural images | 0.0052 | 0.357 | 0.988 | 0.09 |  | 27.9 | 0.22 | 0.052 |
| PLIP | Text anchor (web), from CLIP | 0.0055 | 0.460 | 0.990 | 0.06 |  | 33.8 | 0.38 | 0.042 |

**Figure S5:**
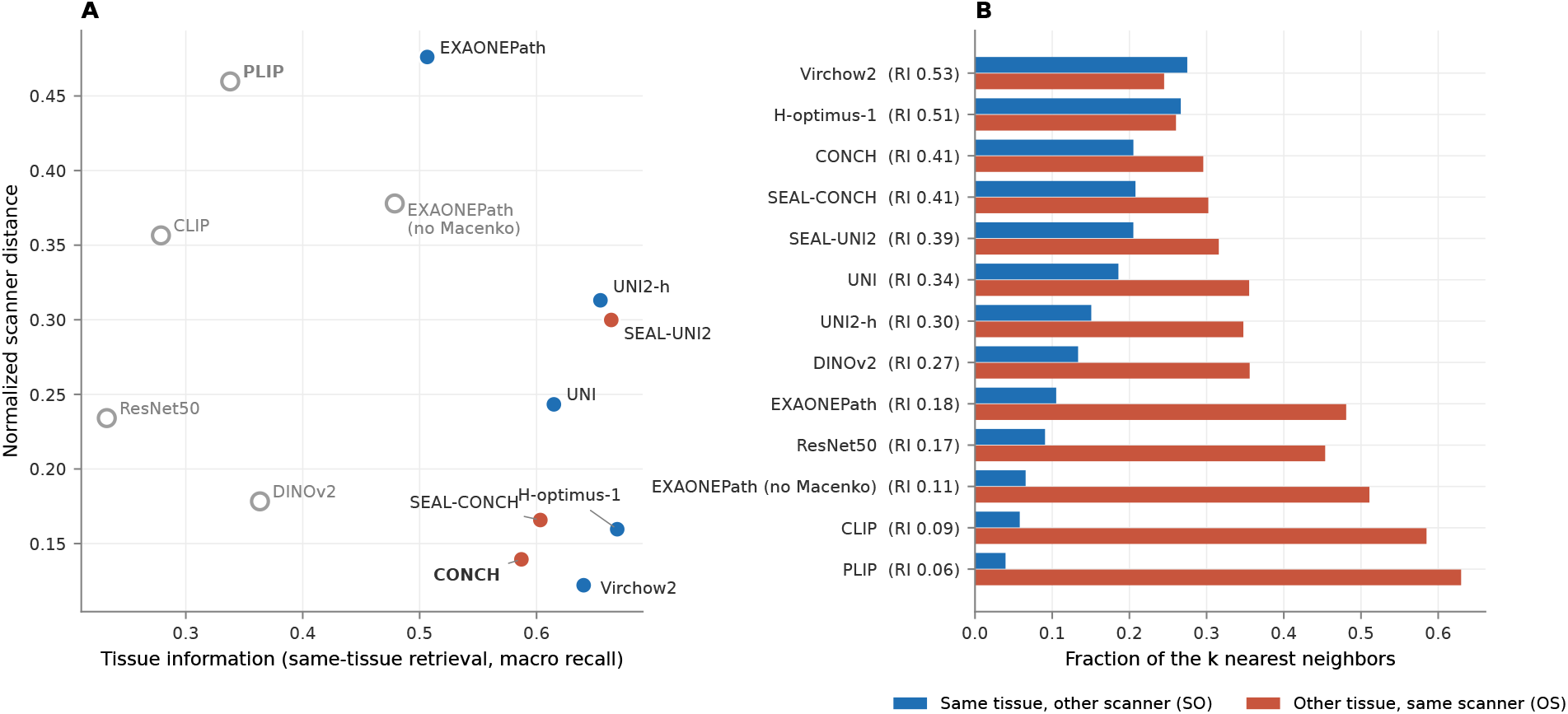
Tissue information and scanner distance for all model variants. **(A)** Same-tissue retrieval against normalized scanner distance. **(B)** Fractions of the *k* nearest neighbors used to compute the robustness index: same tissue from another scanner (SO) and other tissue from the same scanner (OS). Models are ordered by robustness index.

**Figure S6:**
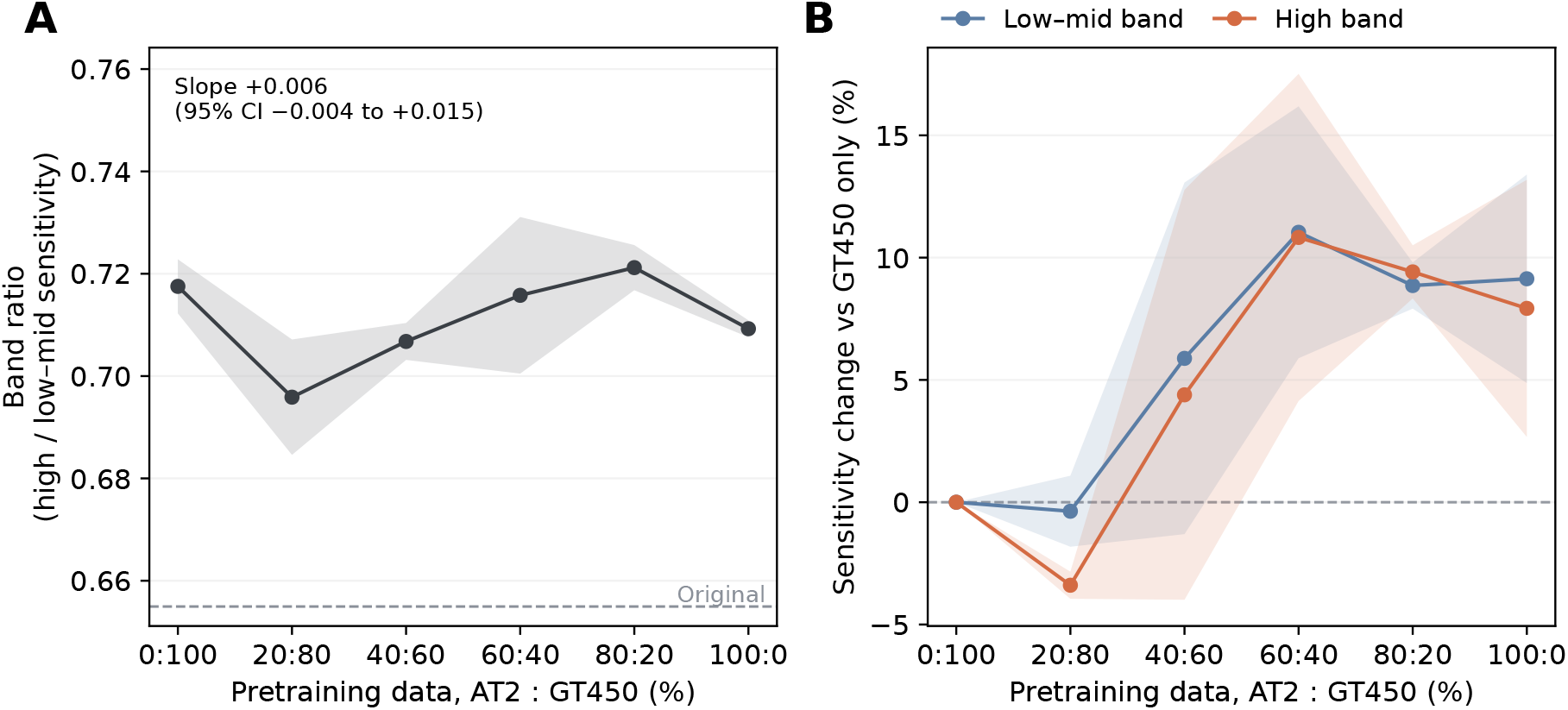
The same experiment in Virchow2, with six mixing proportions. Panels as in Fig. 7.

## Ethics approval

This study was reviewed and deemed exempt by the Institutional Review Board of Children’s Hospital of Philadelphia (IRB 25-024168, “A Multi-Scanner Reference Dataset to Quantify and Mitigate Technical Variation in Digital Pathology”). The requirement for informed consent was waived by the IRB, and all slides were de-identified prior to analysis.

## Data and code availability

PanNormal data are available from the corresponding authors upon request. The PLISM-wsi data used for external evaluation are publicly available on Figshare+ (https://plus.figshare.com/articles/dataset/Pathology_Images_of_Scanners_and_Mobilephones_PLISM_-_Whole_Slide_Images_Dataset/23614422). Analysis code will be made publicly available upon journal submission.

## CRediT authorship contribution statement

**Jonghyun Lee:** Formal analysis, Writing – original draft. **Lina Takemaru:** Formal analysis, Writing – original draft. **Yutong Zhu:** Writing – review & editing. **William Sullivan:** Writing – review & editing. **Matthew E. Lee:** Writing – review & editing. **Angela Ge:** Writing – review & editing. **Portia A. Kreiger:** Resources. **Amy E. Baxter:** Resources. **Zhi Huang:** Writing – review & editing. **Dokyoon Kim:** Supervision. **Derek A. Oldridge:** Supervision.

